# Cross-activation of the FGF, TGF-β and WNT pathways constrains BMP4-mediated induction of the Totipotent state in mouse embryonic stem cells

**DOI:** 10.1101/2022.04.15.488509

**Authors:** Thulaj Meharwade, Loïck Joumier, Maxime Parisotto, Vivian Huynh, Edroaldo Lummertz da Rocha, Mohan Malleshaiah

**Affiliations:** Montreal Clinical Research Institute (IRCM), 110 Pine Avenue West, Montreal, QC H2W 1R7, Canada; Department of Biochemistry and Molecular Medicine, University of Montreal, C.P. 6128, Succursale Centre-ville, Montreal, QC H3C 3J7, Canada; Molecular Biology Program, University of Montreal, C.P. 6128, Succursale Centre-ville, Montreal, QC H3C 3J7, Canada; Department of Microbiology, Immunology and Parasitology, Federal University of Santa Catarina, Florianópolis SC, Brazil; The Division of Experimental Medicine, McGill University, 1001 Decarie Boulevard, Montreal, QC H4A 3J1, Canada; McGill Regenerative Medicine Network, 1160 Pine Avenue West, Montreal, QC H3A 1A3, Canada

**Keywords:** Embryonic stem cells, BMP4 signaling, Totipotent state, signaling cross-activation, stem cell heterogeneity

## Abstract

Cell signaling induced cell fate determination is central to stem cell and developmental biology. Embryonic stem cells (ESC) are an attractive model for understanding the relationship between cell signaling and cell fates. Cultured mouse ESCs can exist in multiple cell states resembling distinct stages of early embryogenesis, such as Totipotent, Pluripotent, Primed and Primitive Endoderm. The signaling mechanisms regulating the Totipotent state acquisition and coexistence of these states are poorly understood. Here we identify BMP4 as an inducer of the Totipotent state. However, we discovered that BMP4-mediated induction of the Totipotent state is constrained by the cross-activation of FGF, TGF-β and WNT pathways. We exploited this finding to enhance the proportion of Totipotent cells in ESCs by rationally inhibiting these cross-activated pathways using small molecules. Single-cell mRNA-sequencing further revealed that induction of the Totipotent state is accompanied by the suppression of both the Primed and Primitive Endoderm states. Furthermore, the reprogrammed Totipotent cells generated in culture have a molecular and functional resemblance to Totipotent cell stages of preimplantation embryos. Our findings reveal a novel BMP4 signaling mechanism in ESCs to regulate multiple cell states, potentially significant for managing stem cell heterogeneity in differentiation and reprogramming.

## INTRODUCTION

Cells use multiple signal transduction pathways to sense and respond to extracellular stimuli. The stimuli information is processed through signaling networks resulting in specific changes in gene expression programs that underly diverse adaptive responses, such as proliferation and differentiation. Understanding how signal transduction mechanisms impact cell fate is fundamental to cell and developmental biology. Even though signaling pathways have been studied extensively, it is not completely resolved how they map to specific downstream cell fate determining gene expression programs. Why do cells often respond to a single stimulus by activating multiple pathways in parallel?

Mouse embryonic stem cells (ESCs), with their ability to self-renew and differentiate into embryonic cell types (referred to as pluripotency), represent a valuable experimental system to study the relationship between cell signaling pathways and gene expression programs of cell fate specification (*1, 2*). Multiple developmental signaling pathways such as BMP, FGF, TGF-β, WNT and LIF regulate ESCs. One or more of these pathways, often in combination, either promote the maintenance of ESCs or induce their differentiation to specific cell fates of early embryogenesis (**Figure 1A**). The LIF pathway promotes the self-renewal of ESCs. It induces the expression of pluripotency genes such as Klf4 to maintain ESCs in the Naive state (pluripotent cells of the preimplantation epiblast) (*3, 4*). The BMP4 pathway promotes both self-renewal and pluripotency by inducing the expression of Id genes to inhibit ESC differentiation (*5*). BMP4 also facilitates the establishment of a stem cell state during the reprogramming of somatic cells to induced pluripotent stem cells (*6*). In combination with LIF, the WNT pathway can promote ESCs by inducing the expression of core pluripotency genes such as Nanog, Sox2 and Oct4 (*7, 8*). However, in the absence of LIF, WNT induces ESC differentiation to the Primitive Endoderm state (extra-embryonic endoderm precursor) (*9*). In contrast to the pluripotency-promoting pathways, the FGF and TGF-β pathways induce mouse ESC differentiation to the Primed state (postimplantation epiblast cells) (*10, 11*). Inhibition of the FGF and TGF-β pathways facilitates the dedifferentiation of Primed cells to Naive state ESCs (*12*). The FGF pathway was also shown to induce the Primitive Endoderm state (*13*). These highlight the critical role of various signaling pathways in regulating ESCs, resulting in multiple culture conditions for their *in vitro* maintenance and propagation (*3, 5, 7, 8*).

**Figure 1:**
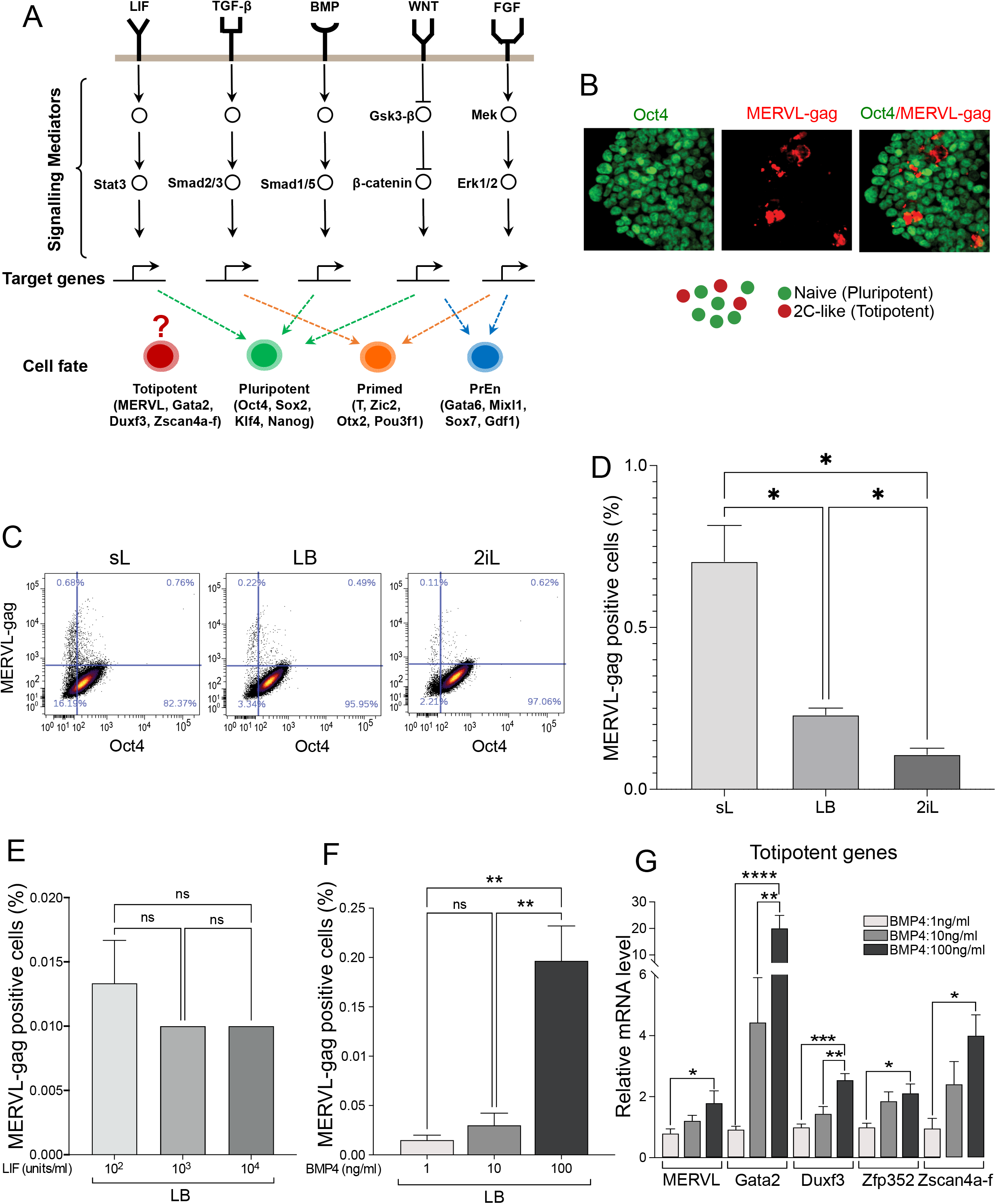
BMP4 induces the Totipotent state in mouse embryonic stem cells. **(A)** Schematic of the common culture condition-related signaling pathways known to promote various cell states in mouse ESCs: Pluripotent (Naive), Totipotent (2C-like), Primed and PrEn (Primitive Endoderm). **(B)** Immunofluorescence images of Oct4 (green) and MERVL-gag (red) stained ESCs cultured in sL (serum + LIF). **(C)** Scatter plots showing flow cytometry analysis of MERVL-gag and Oct4 immunostained ESCs in different culture conditions (sL: serum + LIF; LB: LIF + BMP4; 2iL: Gsk3-β-i + Mek-i + LIF). **(D)** Proportions of MERVL-gag positive cells determined by flow cytometry in ESC culture conditions. Proportions of MERVL-gag positive cells determined by flow cytometry in LB with increasing concentration of LIF **(E)** or BMP4 **(F)**. **(G)** Relative expression of Totipotent genes in ESCs cultured for 72 h in LB with increasing concentration of BMP4. Error bars indicate the mean ± standard error of the mean (SEM) of three (D, E & F) or four (G) biological replicates. ANOVA: ns, nonsignificant; *p≤ 0.05; **p≤ 0.01; ***p≤ 0.001; ****p≤ 0.0001. The displayed images and scatter plots are representative of three independent experiments.

Cultured ESCs exhibit heterogeneous expression of pluripotency genes, including Nanog, Rex1 and Klf4, implying their variable functional potentials (*14*). In addition to the preimplantation epiblast resembling Naive state, ESCs have been reported to exist in other cell states of early embryogenesis. For instance, cultured ESCs can also exist in differentiated states such as Primed and Primitive Endoderm (*15, 16*) (**Figure S1A-B**). Furthermore, ESCs have also been found to acquire a higher level of developmental potency by exhibiting characteristics of totipotency, commonly referred to as the 2C-like state, resembling the 2-Cell stage of preimplantation embryos (*17*) (**Figure 1B**). Unlike Pluripotent cells, Totipotent cells observed in culture can give rise to both embryonic and extra-embryonic cell types (*17, 18*). Such versatile transitions of cultured ESCs to more differentiated and least differentiated states suggest that they might be more plastic in nature than previously thought. For consistency, henceforth, we will refer to various cell states of ESCs as the following: 2C-like cells as “Totipotent”; Naive cells as “Pluripotent”; differentiation-primed cells as “Primed” and Primitive Endoderm cells as “PrEn”.

The cell states observed in ESCs and their counterparts during early embryogenesis are characterized by the distinct expression of their key regulatory genes (**Figure 1A**). The expression of MERVL (murine endogenous retroviral element), Gata2, Duxf3 and Zscan4a-f mark Totipotent cells (*17, 19, 20*). The transcription factor Gata2 induces the Totipotent state by inducing MERVL expression (*19*). The Totipotent state can also be enhanced in cultured ESCs in various ways: over-expression of Duxf3 and Nelfa genes and miR-344 microRNA; altering chromatin states through knockout of Lsd1 and Hdac inhibitors; altering metabolic states through sodium acetate and α-ketoglutarate; treating with a combination of small-molecule inhibitors and retinoic acid (*21–27*). Pluripotent cells are marked by the expression of Oct4, Sox2, Klf4, Nanog, Rex1 and other transcription factors known to form a self-reinforcing gene regulatory network (*28, 29*). Over-expression of Oct4, Sox2, Klf4 and Myc (Yamanaka factors) is sufficient to induce the Pluripotent state in mouse fibroblasts (*30*). Over-expression of Klf4 in Primed cells can also reprogram them to the Pluripotent state (*31*). Primed cells are marked by the expression of Brachyury, Zic3, Otx2 and Pou3f1, and PrEn cells are marked by the expression of Gata6, Mixl1, Sox7 and Gdf1 transcription factors (*32–38*).

The mechanisms regulating ESC’s ability to acquire diverse cell states in the same culture are not fully understood. Moreover, it remains unresolved which signaling pathway(s) among those commonly active in ESC culture conditions is essential to induce the Totipotent state.

In this study, we identified BMP4 signaling to induce the Totipotent state in ESC cultures. However, we found that the BMP4-dependent induction of this state is curtailed by the cross-activation of the FGF, TGF-β and WNT pathways involved in promoting alternative cell states. We could enhance the proportion of Totipotent cells in ESCs and suppress the emergence of Primed and PrEn cells by inhibiting the pathways cross-activated by BMP4. Our findings reveal novel signaling mechanisms regulating the Totipotent state and cell state heterogeneity in ESCs.

## RESULTS

### BMP4 signaling induces the Totipotent state in ESCs

In mouse ESC cultures, a minor proportion of Totipotent cells are observed (*17*) (**Figure 1B**). To determine how these cells arise, we first sought to determine how culture conditions influence the proportion of Totipotent cells. We compared mouse ESCs cultured in three common conditions consisting of basal medium supplemented with either sL (serum and LIF), LB (LIF and BMP4) or 2iL (LIF, CHIR99021 (GSK3-β inhibitor) and PD0325901 (MEK inhibitor)) (*3, 5, 7, 8*). We analyzed proportions of Totipotent and Pluripotent cells using flow cytometry by performing immunostaining of their previously reported cell state marker proteins, MERVL-Gag (MERVL) and Oct4, respectively (*17, 29*) (**Figure 1C**). Although the overall proportion of Totipotent cells is less than 1%, the highest proportion was observed in sL (~0.7%), followed by LB (~0.2%) and 2iL (~0.1%) (**Figure 1D**). These data suggest that: 1) signals within sL and LB may promote the Totipotent state, but such signals might either be lacking or inhibited in 2iL, a well-known condition to promote the Pluripotent state robustly (*8*); 2) variations in cell state proportions are unlikely to be induced by LIF since it is common to all conditions; 3) BMP4 in LB, and sL (serum is also a source of BMP4 (*39*)) may induce the Totipotent state in ESCs.

To test the potential role of BMP4 and LIF in inducing the Totipotent state, we individually varied their concentrations in LB. Varying LIF concentration 10-fold less or more relative to its regular concentration (1000 units/ml) did not induce significant changes in the proportion of Totipotent cells (**Figure 1E**). On the other hand, increasing BMP4 concentration by 10-fold (100 ng/ml) from its regular concentration (10ng/ml) resulted in a 5- to 10-fold increase in the proportion of Totipotent cells (**Figure 1F**). We further tested the effect of BMP4 on the Totipotent state by assessing the expression of multiple marker genes using quantitative PCR (qPCR). We observed an increase in the expression of MERVL, Gata2, Duxf3, Zfp352 and Zscan4a-f with an increased BMP4 concentration in LB (**Figure 1G**). Since BMP4 is known to promote ESCs by blocking their differentiation (*5*), we analyzed the expression of Pluripotent genes Oct4, Nanog, Sox2, Klf4 and Rex1. While Oct4, Nanog and Sox2 did not change significantly, Klf4 and Rex1 expression showed a concentration-dependent increase with BMP4 (**Figure S1C**).

In addition to the Totipotent state, Primed and PrEn cell state signatures have also been observed in cultured ESCs (*15, 16*) (**Figure S1A-B**). Thus, we wondered whether BMP4 influences these alternative cell states. To test this, we measured the expression pattern of Primed and PrEn genes in LB with varying concentrations of BMP4. Indeed, we observed a BMP4 concentration-dependent increase in the expression of Primed genes: Otx2, Zic3 and Brachyury (**Figure S1D**), and PrEn genes: Gata6, Mixl1 and Sox7 (**Figure S1E**).

These data altogether indicate that BMP4 signaling induces the Totipotent, Primed and PrEn states, in addition to its known pluripotency-promoting role in ESCs. How might the BMP4 signaling pathway induce multiple distinct cell states in ESCs?

### BMP4 cross-activates the FGF, TGF-β and WNT pathways

Since BMP4 induced the Totipotent state, as well as Pluripotent, Primed and PrEn states which are known to be induced by either WNT, TGF-β or FGF signaling (**Figure 1A**), we hypothesized that BMP4 cross-activates these pathways to promote multiple cell states in ESCs.

To determine whether BMP4 cross-activates the FGF, TGF-β and WNT pathways, we first assessed the change in activation levels of their critical signaling mediator proteins in a BMP4 concentrationdependent manner. We evaluated the activation of each pathway based on the levels of pSMAD1/5 (BMP4), pERK1/2 (FGF), pSMAD2/3 (TGF-β), and total β-Catenin (WNT) (**Figure 1A**). To perform these experiments, ESCs were pre-seeded in sL and switched to LB with increasing concentrations of BMP4 (1 to 100 ng/ml) for 24 h. We observed an apparent BMP4 concentration-dependent increase in pSMAD1/5 levels (**Figure 2A**). In addition, we observed a similar trend in pERK1/2 and pSMAD2/3 levels, saturated at 100 ng/ml of BMP4. Total β-Catenin levels peaked at 1 to 5 ng/ml of BMP4 and decreased at higher concentrations. Thus, these data demonstrate that BMP4 signaling also activates the FGF, TGF-β and WNT pathways.

**Figure 2:**
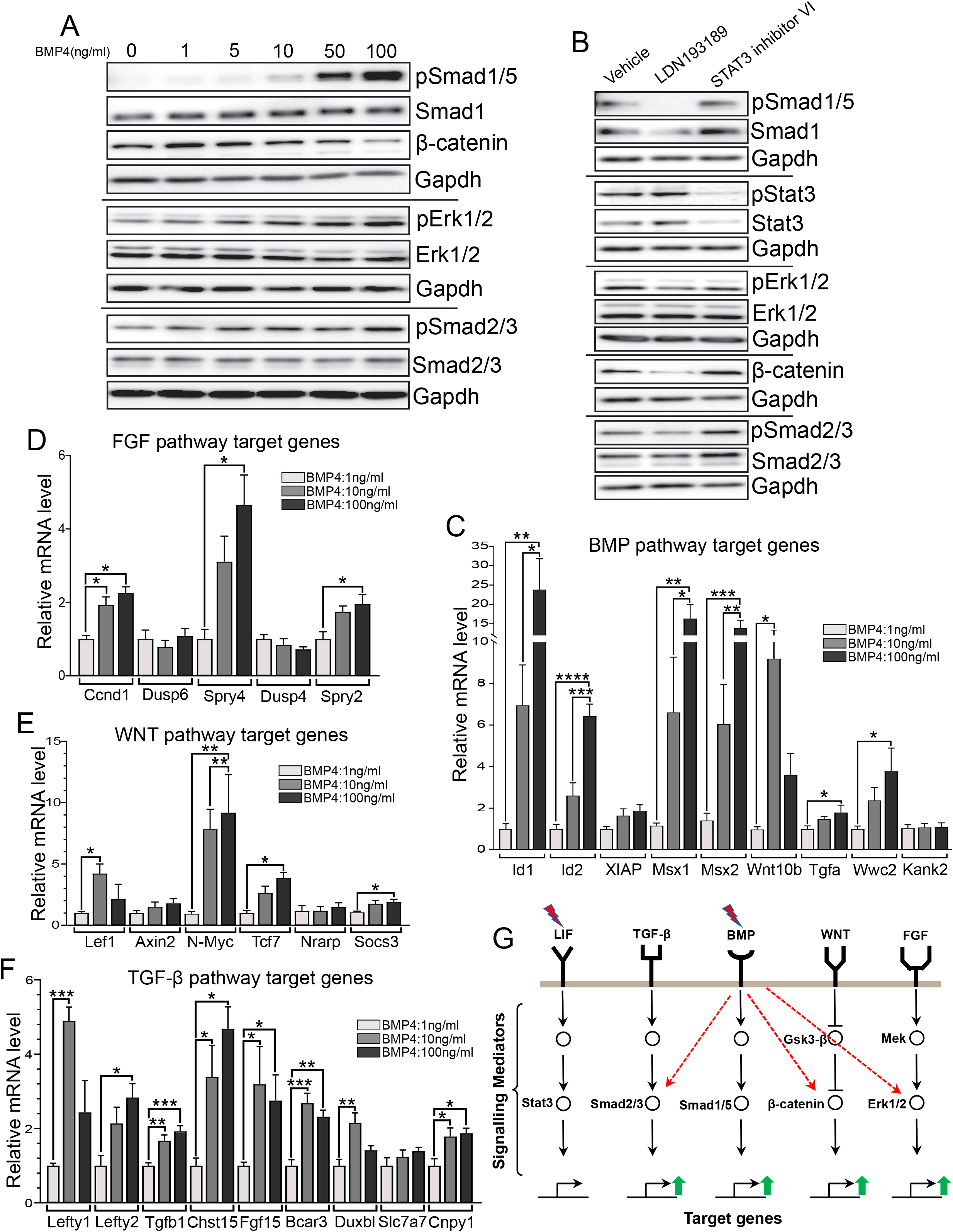
BMP4 signaling cross-activates the FGF, TGF-β and WNT pathways. **(A)** Western blot analysis of BMP4, WNT, FGF, and TGF-β signaling pathways in ESCs cultured in LB with increasing concentrations of BMP4. **(B)** Western blot analysis of BMP4, LIF, FGF, WNT and TGF-β signaling pathway activation in ESCs cultured in LB for 24 h in the presence of either a BMP4 pathway activation inhibitor (LDN193189) or Stat3 inhibitor (Stat3 inhibitor VI) or vehicle. Gapdh was used as loading control in A and B. Displayed images are representative of four (A) or three (B) independent experiments. Relative expression of BMP4 pathway target genes **(C)**, FGF pathway target genes **(D)**, WNT pathway target genes **(E)** and TGF-β pathway target genes **(F)** in ESCs grown in LB for 72 h with increasing concentration of BMP4. **(G)** Schematic showing BMP4-mediated cross-activation of the FGF, WNT and TGF-β pathways in mouse ESCs. Error bars indicate the mean ± SEM of three biological replicates. ANOVA: *p≤ 0.05; **p≤ 0.01; ***p≤ 0.001; ****p≤ 0.0001.

To confirm that the cross-activations observed in LB were indeed due to BMP4 signaling and not LIF, we blocked their activation with specific pharmacological inhibitors. We used LDN193189, a selective inhibitor of BMP type I receptors ALK1/2/3, to prevent the activation of BMP4 signaling, and STAT3 Inhibitor VI to prevent LIF signaling (*40, 41*). As expected, activation of BMP4 signaling (pSMAD1/5) and LIF signaling (pSTAT3) were abolished in ESCs treated with LDN193189 and STAT3 Inhibitor VI, respectively (**Figure 2B**). In addition, blocking BMP4 signaling resulted in a drastic reduction in the activation of the FGF (pERK1/2), WNT (total β-Catenin) and TGF-β (pSMAD2/3) pathways. However, preventing LIF signaling did not have a noticeable effect on activating the FGF, TGF-β or WNT pathways (**Figure 2B**). Together, these data confirm that the activation of the FGF, TGF-β and WNT pathways observed in LB is due to BMP4 signaling.

Cell signaling-driven transcriptional responses often induce the expression of genes involved in feedback and autocrine/paracrine signaling mechanisms. We tested the general effect of gene expression on BMP4-dependent activation of pathways by blocking the translation of mRNA to protein using cycloheximide. We observed a rapid activation of the BMP4, WNT, FGF and TGF-β pathways within the first 15 to 30 min of LB addition (**Figure S1F-H**). However, they did not sustain their peak activation for a longer duration. These data suggest that the initial BMP4-dependent cross-activation of the FGF, TGF-β and WNT pathways might be mediated through intracellular signaling and that the sustenance of their activation might require the central dogma.

Next, we asked whether the BMP4-mediated cross-activation is spurious or results in downstream transcriptional responses. We tested this by measuring the BMP4 concentration-dependent changes in the expression of well-described pathway-specific target genes (*5, 42–48*). In agreement with the pathway activation, the expression of known BMP4 pathway target genes increased in a concentration-dependent manner (**Figure 2C**). Multiple known target genes of the FGF, WNT, and TGF-β pathways also displayed a similar trend (**Figure 2D-F**). Induction of pathway target gene expression agrees with the activation of these pathways by BMP4 (**Figure 2A-B**). Together, these data indicate that BMP4-mediated cross-activation of multiple pathways is not just spurious but results in pathway-specific downstream responses (**Figure 2G**).

### Inhibition of the BMP4-mediated activation of FGF, TGF-β and WNT pathways enhances the Totipotent state in ESCs

Our results indicate that BMP4 induces the Totipotent state in ESCs, but within a constrained capacity as it also cross-activated the FGF, TGF-β and WNT pathways required to promote alternative cell states. Thus, we hypothesized that inhibiting the BMP4 cross-activated pathways may enhance its ability to induce the Totipotent state while restricting it from inducing alternative cell states. To test this, we rationally inhibited the cross-activated pathways using small molecules and assessed their effect on inducing the Totipotent state (**Figures 3A & S2A**). Since the BMP4-mediated cross-activations were observed in LB (**Figure 2**), we used it as a basal condition for our tests. We used flow cytometry to measure the proportion of Totipotent cells (MERVL+) in ESCs treated with various combinations of ligands and inhibitors for 72 h (**Figure 3B**). As observed before in common culture conditions, the proportion of Totipotent cells decreased from sL, LB to 2iL (**Figures 1D & 3B**).

**Figure 3:**
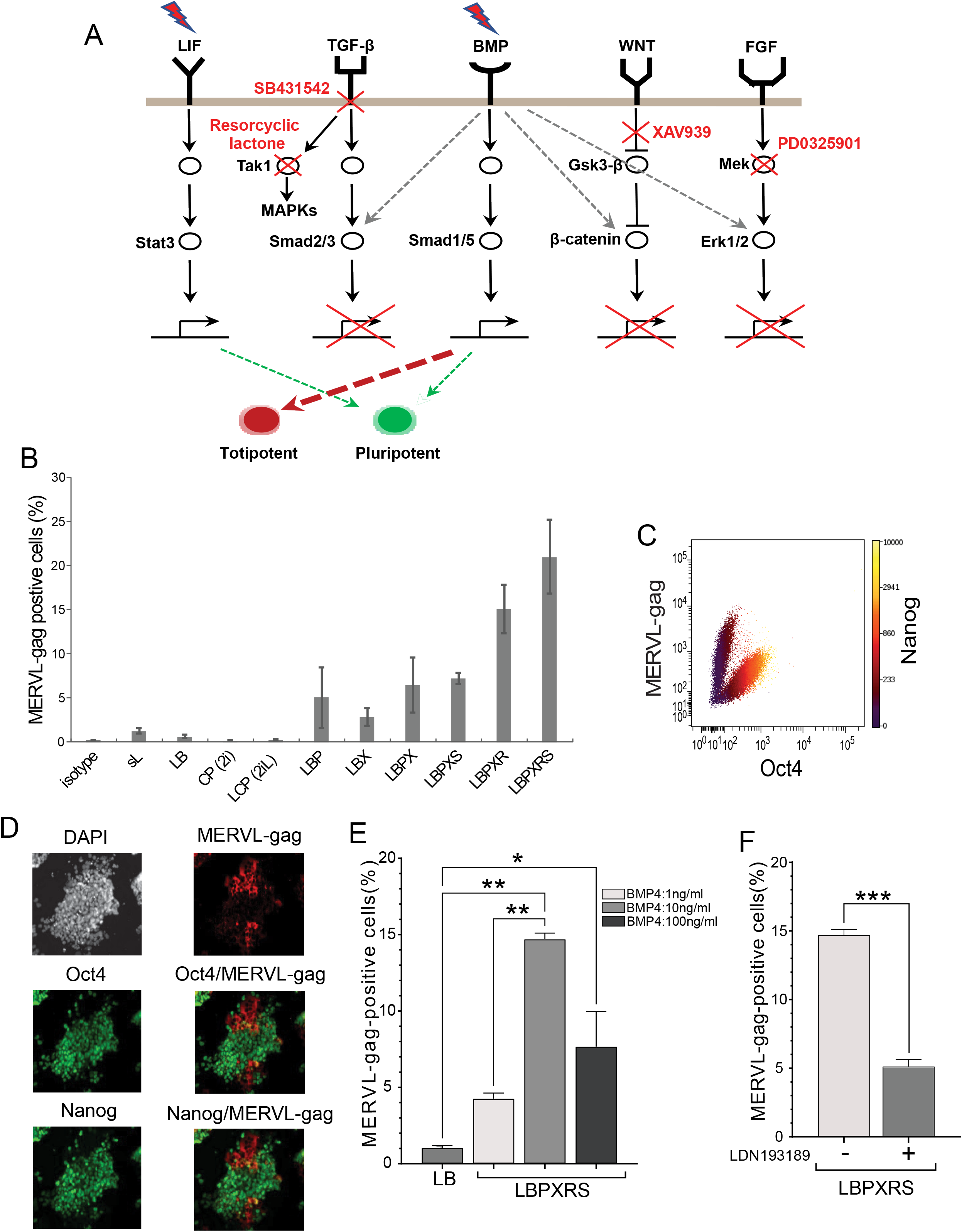
Inhibition of the BMP4-mediated activation of FGF, TGF-β and WNT pathways enhances Totipotent cell proportions in ESCs. **(A)** Schematic showing the rational strategy used to inhibit BMP4 cross-activated FGF, TGF-β and WNT pathways using combinations of ligands and small molecule inhibitors. Also, see Figure S2A for specific details of each combination. **(B)** The proportion of MERVL-gag positive cells determined by flow cytometry analysis of immunostained ESCs cultured for 72 h in the indicated conditions. s: serum, L: LIF, B: BMP4, C: CHIR99021, P: PD0325901, X: XAV939, S: SB431542, R: Resorcyclic lactone. **(C)** Scatter plot showing flow cytometry determined MERVL-gag, Oct4 and Nanog levels in immunostained ESCs cultured for 72 h in LBPXRS. **(D)** Immunofluorescence images of Oct4, Nanog and MERVL-gag stained ESCs cultured in LBPXRS for 72 h. **(E)** The proportion of MERVL-gag positive cells determined by flow cytometry analysis of immunostained ESCs cultured in LB or LBPXRS. ANOVA: *p≤ 0.05, **p≤ 0.01. **(F)** The proportion of MERVL-gag positive cells determined by flow cytometry analysis of immunostained ESCs cultured in LBPXRS in the absence or presence of LDN193189. t-test: ***p≤ 0.001. The displayed images and scatter plots are representative of three independent experiments. Error bars indicate the mean ± SEM of three biological replicates.

To inhibit the cross-activated pathways in LB, we first blocked the FGF pathway (ERK1/2 activation) by using the MEK1/2 inhibitor PD0325901 (forming LBP) (*8, 49*). Interestingly, LBP increased the proportion of Totipotent cells to ~5% (**Figure 3A–B**). Then, we blocked the WNT pathway (β-Catenin accumulation) by using XAV939 in LB (LBX). XAV939 is an inhibitor of Tankyrase 1/2, resulting in stabilized Axin levels required for β-Catenin degradation (*50*). LBX resulted in a modest but significant increase in the proportion of Totipotent cells (~3%) (**Figure 3B**). In addition, inhibition of both the FGF and WNT pathways with PD0325901 and XAV939, respectively, in LB (LBPX) increased the proportion of Totipotent cells (~6%).

Next, we inhibited the TGF-β pathway by using SB431542 in LBPX (forming LBPXS). SB431542 is a well-described TGF-β pathway-specific inhibitor that does not inhibit the related BMP4 Type I receptors (*51*). LBPXS resulted in ~8% of Totipotent cells (**Figure 3B**). TGF-β signaling can also activate multiple MAPKs such as p38, JNK and ERK1/2 through TAK1 (TGF-β-activated kinase 1) (*52*). We, therefore, used Resorcyclic lactone, a specific inhibitor of TAK1, to prevent the potential activation of multiple MAPKs through TGF-β signaling (*53*). Resorcyclic lactone in LBPX (LBPXR) increased the proportion of Totipotent cells to ~15%. Finally, inhibiting all three pathways using PD0325901, XAV939, Resorcyclic lactone and SB431542 in LB (LBPXRS) enhanced the proportion of Totipotent cells to ~20% (**Figure 3B**). LBPXRS increased the proportion of Totipotent cells by ~105-fold relative to 2iL (a robust ESC culture condition), ~35-fold relative to LB and ~4-fold relative to LBP (**Figure 3B**). In addition, Totipotent cells in LBPXRS showed reduced levels of Pluripotent state genes Oct4 and Nanog (**Figure 3C-D**).

To confirm whether the increase in the proportion of Totipotent cells observed in LBPXRS is indeed due to BMP4 signaling, we varied the concentration of BMP4 in LBPXRS and blocked BMP4 pathway activation using LDN193189. Decreasing BMP4 by 10-fold in LBPXRS drastically reduced the proportion of Totipotent cells (**Figure 3E**). Intriguingly, increasing BMP4 by 10-fold in LBPXRS also resulted in a reduction of Totipotent cells, suggesting that the hyperactivation of BMP4 signaling beyond a certain level might compromise its function. The addition of LDN193189 prevented the increase in the proportion of Totipotent cells observed in LBPXRS (**Figure 3F**). Together, these data confirm the essential role of BMP4 signaling in inducing the Totipotent state in ESCs.

Next, we asked whether cross-activation of the FGF, TGF-β and WNT pathways were indeed prevented by the combination of inhibitors used in LBPXRS. To determine this, we measured the changes in activation levels of critical signaling mediator proteins in LB and LBPXRS with increasing concentrations of BMP4 (0 to 100 ng/ml). Surprisingly, as reflected by pSMAD1/5 levels, we observed a 2- to 3-fold higher activation of BMP4 signaling in LBPXRS compared to LB (**Figure 4A-B**). On the contrary, as reflected by pERK1/2 and pSMAD2/3 levels, activation of the FGF and TGF-β pathways was abolished in LBPXRS (**Figure 4C-D**). However, we failed to observe a significant decrease in β-Catenin levels in LBPXRS, suggesting an alternative mechanism of its inhibition by XAV939 (**Figure 4A**). It has been reported that XAV939 can inhibit β-Catenin by preventing its cytosol-to-nuclear translocation (*54, 55*). To test whether this mechanism operates in LB vs LBPXRS, we performed immunostaining of β-Catenin and DAPI staining of nuclear DNA, followed by imaging and quantification of the nuclear fraction of β-Catenin (**Figure S2B**). As expected, WNT pathway activation using Wnt3a resulted in an increased fraction of cells with nuclear β-Catenin (60% of the nuclei) (**Figure S2C**). While this fraction remained relatively high in LB (45% of the nuclei), it decreased to ~10% in LBPXRS. In agreement with decreased nuclear translocation of β-Catenin, we observed its increased accumulation at the membrane/cytosol in LBPXRS compared to LB (**Figure S2B**). Thus, these data indicate that XAV939 in LBPXRS inhibits β-Catenin by disrupting its nuclear translocation but not affecting its protein levels.

**Figure 4:**
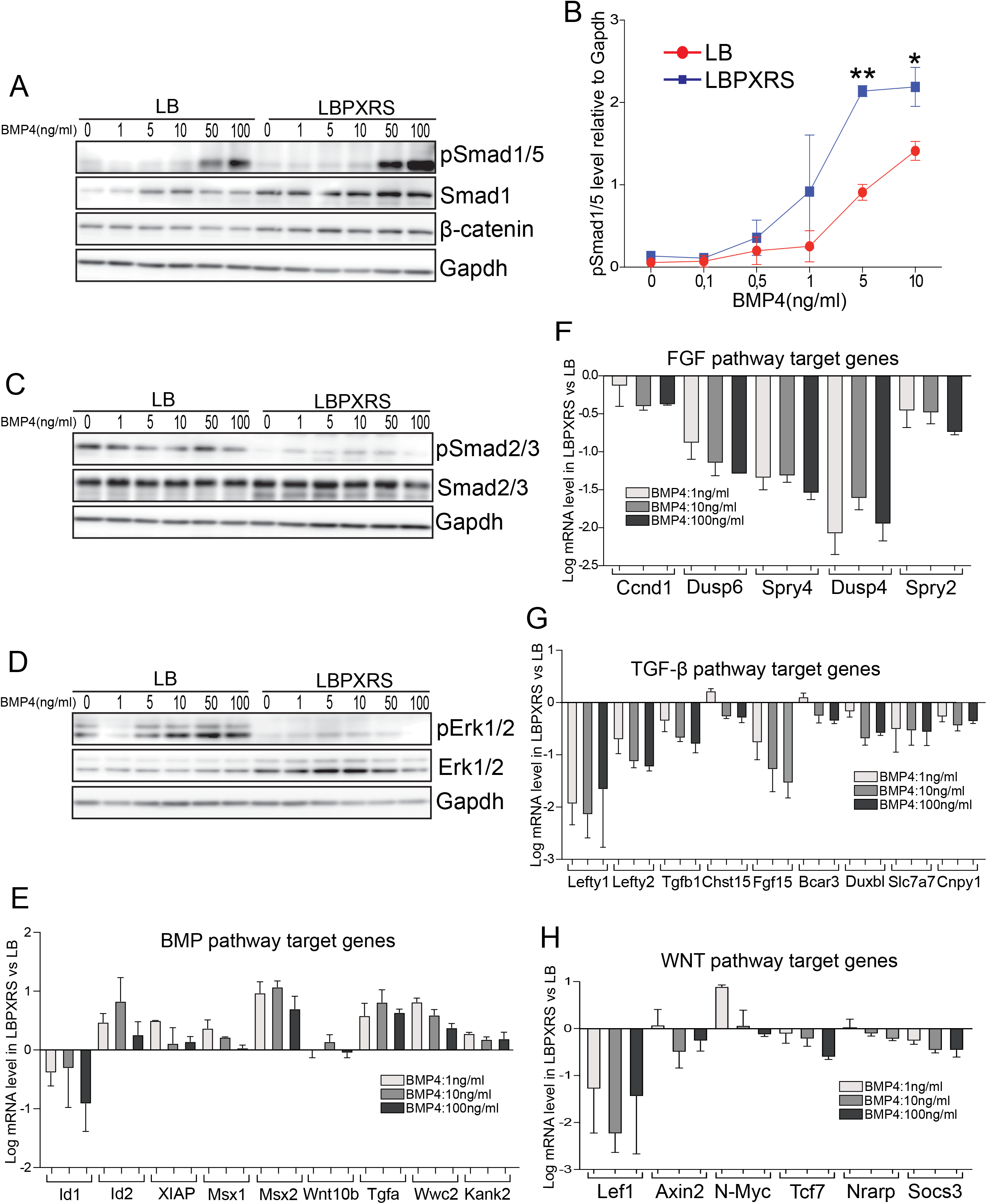
Inhibition of the BMP4 cross-activated pathways enhances the intensity of BMP4 signaling. **(A)** Western blot analysis of BMP4 and WNT signaling pathways in ESCs cultured in LB or LBPXRS with increasing concentrations of BMP4. **(B)** Quantification of relative pSmad1/5 levels from western blots of ESCs cultured in LB or LBPXRS with increasing concentrations of BMP4. Error bars indicate the mean ± SEM of three biological replicates. t-test: *p≤ 0.05; **p≤ 0.01. Western blot analysis of TGF-β **(C)** and FGF **(D)** signaling pathways in ESCs cultured in LB or LBPXRS with increasing concentrations of BMP4. In A, C & D; Gapdh was used as a loading control, and the displayed images are representative of three independent experiments. Relative expression (log scale, relative to LB) of the indicated target genes of the BMP4 **(E)**, FGF **(F)**, TGF-β **(G)** and WNT **(H)** pathways in cells cultured in LBPXRS or LB for 72 h with increasing concentrations of BMP4. Error bars indicate the mean ± SEM of four to eight biological replicates.

To determine whether inhibition of the cross-activated pathways also translates to the inhibition of the downstream transcriptional response, we performed the qPCR analysis of canonical target genes of the respective pathways. Relative to LB and in agreement with its enhanced signaling (**Figure 4A-B**), the expression of BMP4 pathway target genes (e.g. Id2, Msx2, Tgfa, Wwc2 and Kank2) was increased in LBPXRS (**Figure 4E**). Also, in agreement with the inhibition of their signaling, expression of FGF, TGF-β and WNT pathway target genes was suppressed in LBPXRS (**Figure 4F-H**). The suppression of WNT pathway target genes in LBPXRS further confirms that inhibiting β-Catenin nuclear translocation with XAV939 inhibits this pathway’s transcriptional response.

Together, these data demonstrate that the induction of the Totipotent state in ESCs is enhanced in accordance with enhanced signaling of the BMP4 pathway and restriction of the cross-activated FGF, TGF-β and WNT pathways.

### Single-cell mRNA-sequencing (scRNA-seq) reveals an enhanced Totipotent state and reduced cell state heterogeneity of ESCs in LBPXRS

The small molecules used were highly effective in inhibiting the BMP4 cross-activated FGF and TGF-β pathways, which are known to induce Primed and PrEn states (*10, 11, 13*). In addition to enhancing the Totipotent state, inhibiting these pathways should reduce the induction of Primed and PrEn states and result in less ESC heterogeneity. To better resolve the effects on multiple cell states simultaneously, we performed scRNA-seq of ESCs cultured in LB and LBPXRS. After standard sequencing data preprocessing, we recovered 1636 cells for LB and 2566 cells for LBPXRS. We integrated both LB and LBPXRS data using Seurat (*56*) to understand their differences better, which showed distinct and similar clusters of cells from both conditions (**Figure S3A**). To classify the cells to their likely cell states, we relied on the relative expression of multiple cell state genes previously described in the literature (**Figures S3B-C & S4**). For instance: Gata2, Tmem191c, Amhr2, Rell1, Aqp3, Trak1, Slc26a11 and Trpm1 regulate the Totipotent state (*18, 19, 57, 58*); Sox2, Klf4, Nanog and Rex1/Zfp42 regulate the Pluripotent state (*28, 29, 32*); Zic3, Brachyury and Fgf8 regulate the Primed state (*32–34*); and Gata6, Cdh2, Mixl1 and Dab2 regulate the PrEn state during early embryo development (*35–38*). Confirming the classification, the annotated cell types maintained exclusive expression patterns of their marker genes (**Figure 5A-B**).

**Figure 5:**
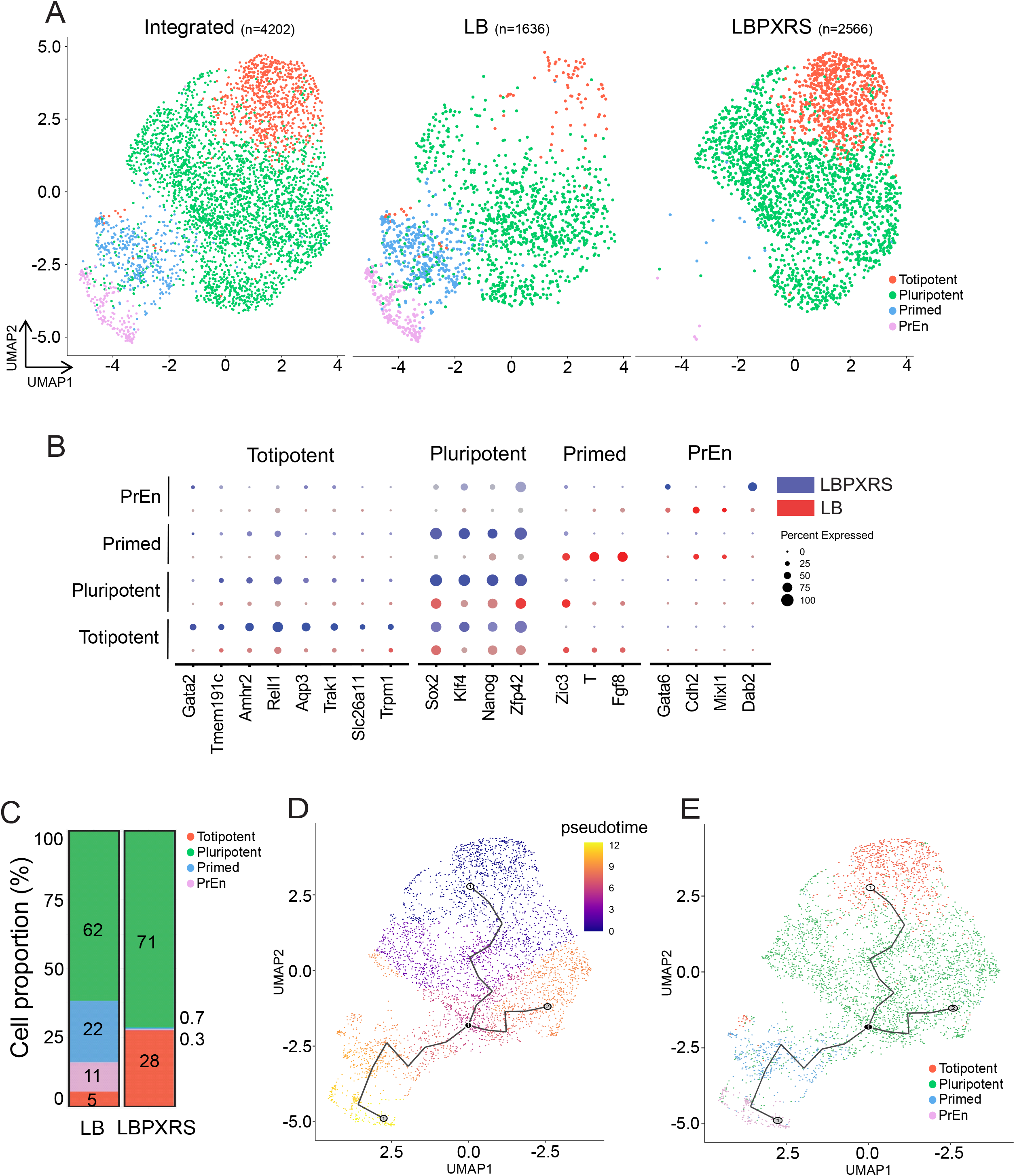
Single-cell mRNA-sequencing reveals an enhanced Totipotent state and a reduced cell state heterogeneity of ESCs in LBPXRS. **(A)** UMAP space of integrated scRNA-seq datasets representing mouse ESCs cultured in LB and LBPXRS conditions. See Figures S3 & S4 and methods for details on annotating the clusters to the indicated cell types (i.e. Totipotent, Pluripotent, Primed and PrEn). **(B)** Relative expression of annotated cell type marker genes in the integrated scRNA-seq dataset of ESCs cultured in LB and LBPXRS. **(C)** Calculated proportions of Totipotent, Pluripotent, Primed and PrEn cells from the integrated dataset. UMAP space showing pseudotime inference and branch analysis **(D)** and cell state membership **(E)**. Totipotent cells were chosen as the source of the pseudotime (i.e. pseudotime = 0).

In LB, all four cell types were detected, with a majority of cells in the Pluripotent state (62%), followed by Primed (22%), PrEn (11%) and a minor proportion of cells in the Totipotent state (5%) (**Figure 5C**). In sharp contrast, LBPXRS mainly consisted of cells in the Pluripotent (71%) and Totipotent (28%) states, with a minimal proportion of cells in the Primed (0.7%) and PrEn (0.3%) states. While increasing the proportion of Totipotent cells by >5-fold, the LBPXRS condition nearly abolished the proportion of both Primed and PrEn cells. In agreement, the absolute mean expression levels of Totipotent and Pluripotent genes were induced in LBPXRS, and Primed and PrEn genes were suppressed (**Figure S5A**). While multiple signaling genes were expressed in LB, they were generally suppressed in LBPXRS (**Figure S5A-B**). We validated the scRNA-seq results for key cell state genes using qPCR analysis in LB and LBPXRS (**Figure S5C-F**). Relative to their expression in LB, MERVL and Totipotent genes (Gata2, Duxf3 and Zfp352) were induced in LBPXRS, whereas both Primed and PrEn genes were suppressed. The expression of multiple Pluripotent genes did not change significantly, except for Oct4, which was induced in LBPXRS (**Figure S5D**). We observed a down-regulation in Zscan4a-f expression in LBPXRS (**Figure S5C**). While Zscan4 is commonly used to assess the Totipotent state, recent evidence suggests that its expression marks an intermediate state between the Pluripotent and Totipotent state (*59*). Suppression of Zscan4 in LBPXRS suggests a plausible alternative mechanism for the Pluripotent-to-Totipotent state transition that might be independent of Zscan4.

scRNA-seq data further confirm the enhanced proportion of Totipotent cells in ESCs cultured in LBPXRS. It also revealed that the BMP4-mediated cross-activation of FGF, TGF-β and WNT pathways results in pronounced cell state heterogeneity in ESCs and that its extent can be reduced by inhibiting these cross-activated pathways.

### Totipotent cells in LBPXRS harbor bidirectional developmental potential

The ordered spatial segregation of cell states in the scRNA-seq data prompted us to evaluate whether the gene expression changes are consistent with the transition of cell states during early embryonic development (**Figure 5A**). We performed a trajectory analysis using Monocle 3 to test this (*60*). The pseudotime trajectory traversed from the source (Totipotent cells) to Pluripotent, followed by Primed and PrEn cells (**Figure 5D-E**). This trajectory path roughly resembles the transition of Totipotent cells into Pluripotent (preimplantation epiblast) and PrEn, followed by the Primed (postimplantation epiblast) state during early embryogenesis (*61*).

Next, we explored the similarity between Totipotent cells in LBPXRS and Totipotent cell stages of the embryo, *in vitro* generated expanded pluripotent stem cells (EPSCs), and other cells from early embryos and cultures. In general, the integration of datasets showed spatially segregated cell stages according to developmental transitions from the Zygote to the E4.5 stage (**Figure 6A**). The relative location of Totipotent cells from LBPXRS suggests that their gene expression pattern is more similar to the embryonic 2-Cell, 4-Cell, 8-Cell cell stages and EPSC-1 (**Figure 6A**). We further verified the similarity in gene expression through hierarchical clustering. Totipotent cells from LBPXRS were classified within the unique branch of totipotency-associated cell stages with a high correlation to EPSC-1, 4-Cell and 8-Cell, followed by 2-Cell and 1-Cell stages (**Figure 6B**). Furthermore, their gene expression was distinct from non-Totipotent cells: morula, E3.0-3.5, E4.5, 2i and Primed.

**Figure 6:**
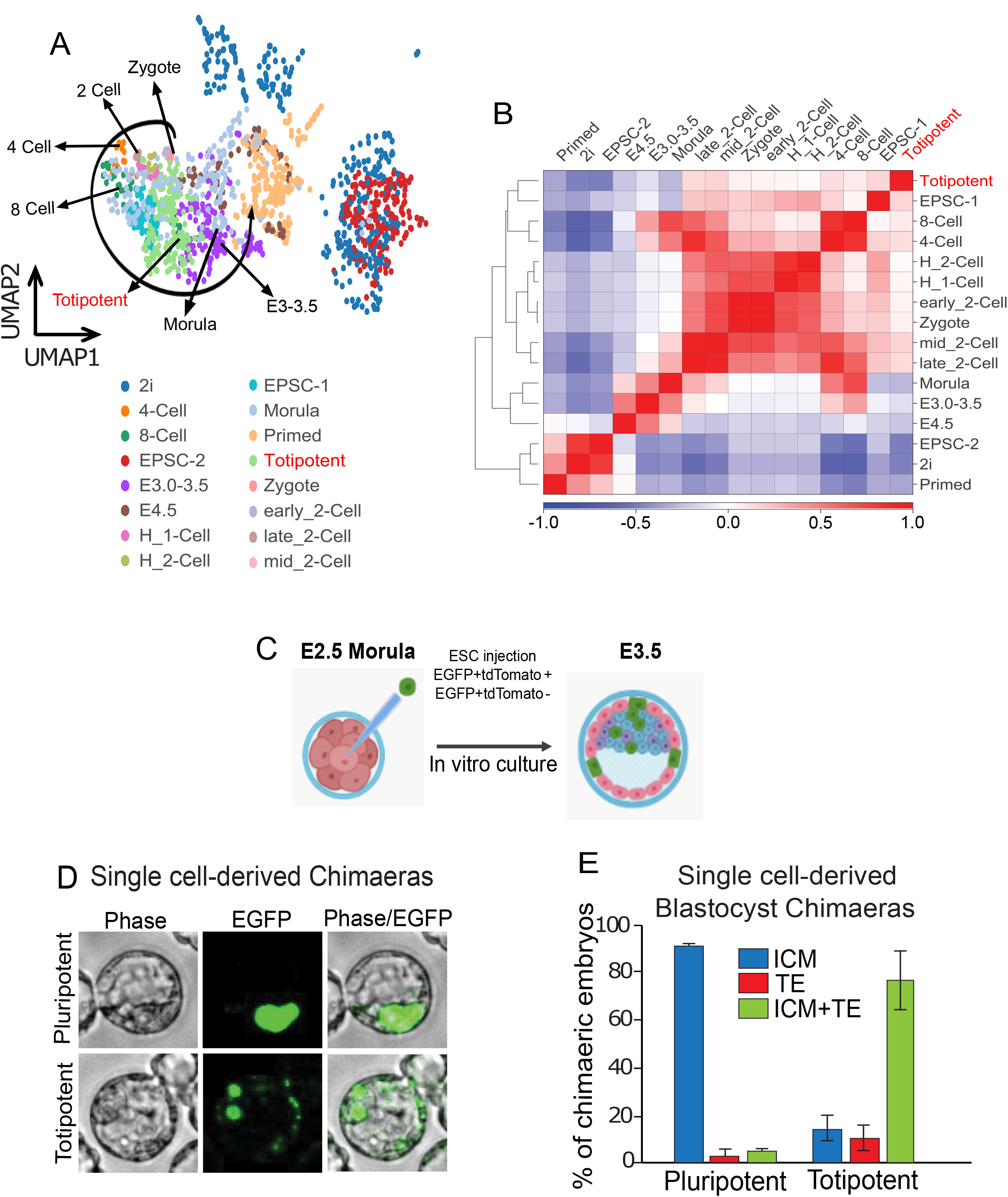
Totipotent cells in LBPXRS harbor bidirectional developmental potential. **(A)** Integration analysis of Totipotent cells in LBPXRS with cells from different cell stages of early embryognesis and ESC cultures and other *in vitro* generated Totipotent-like cells. Manually drawn trajectory showing segregation of early embryogenesis cell stages from Zygote-to-E4.5. **(B)** Hierarchical correlation analysis of the multiple datasets integrated in A. **(C)** Schematic showing the blastocyst chimera generation assay procedure using microinjection of EGFP-labeled cells into the morula of the preimplantation embryo. The microinjected morulae were *in vitro* cultured until the E3.5 stage for blastocyst chimera evaluation. See text and methods for additional details. **(D)** Images of E3.5 blastocysts showing the viability, proliferation and distinct contribution of LBPXRS-cultured Pluripotent cells to the ICM (inner cell mass) (top) and Totipotent cells to both the ICM and TE (trophectoderm) (bottom). MERVL-tdTomato ESCs were stably transfected with an EGFP construct under a constitutively active EF1α promoter to aid cell tracking. **(E)** Proportions of chimeric embryos derived from single Pluripotent or Totipotent cells and their distinctive contributions. A total of 87 Pluripotent (n1 = 40 & n2 = 47) cells and 98 Totipotent cells (n1 = 48 & n2 = 58) were microinjected and recovered blastocyst embryos were analyzed. Error bars indicate standard deviations.

The high gene expression similarity between Totipotent cells from LBPXRS and embryonic cells at Totipotent stages of development suggests their functional resemblance. Totipotent cells are characterized by their ability to give rise to both the inner cell mass (ICM) and trophectoderm (TE), referred to as bidirectional developmental potential (*62*). To determine whether Totipotent cells in LBPXRS are bidirectional, we assessed their ability to form chimeric blastocysts and E12.5 embryos following their microinjection at the morula stage (**Figures 6C & S6**). We used 2-Cell reporter cells (MERVL::tdTomato) stably transfected with an EF1α-driven EGFP-expressing plasmid construct to mark the progeny of microinjected cells (*17*). Single, sorted EGFP+ MERVL::tdTomato+ and EGFP+ MERVL::tdTomato-cells cultured in LBPXRS were microinjected into embryos at the morula stage (E2.5) and allowed to develop into blastocysts (E3.5). The microinjected single cells integrated within the embryos (**Figure 6D**). In blastocyst chimeras, tdTomato-cells primarily contributed to the ICM and tdTomato+ cells contributed efficiently to both the ICM and TE (**Figure 6E**). In E12.5 chimeric embryos, tdTomato-cells only contributed to embryonic tissues (mesoderm, endoderm, ectoderm) and tdTomato+ cells contributed to both embryonic and extra-embryonic tissues (placenta and yolk sac) (**Figure S6**). Similarly, tissue-integrated tdTomato-cells co-expressed only embryonic tissue markers (e.g. Brachyury and Sox1), and tdTomato+ cells co-expressed both embryonic as well as extra-embryonic tissue markers (e.g. Cdx2 and Tpbpa) (**Figure S7**).

Taken together, scRNA-seq and microinjection results demonstrate that Totipotent cells in LBPXRS are similar to Totipotent cells of the preimplantation embryo at both the molecular and functional levels.

## DISCUSSION

In this study, we investigated signaling mechanisms that regulated the Totipotent state in ESCs and identified the essential roles of BMP4. In addition to inducing the Totipotent state, we discovered that BMP4 signaling promotes other cell states of ESCs such as Pluripotent, Primed and PrEn by cross-activating the FGF, TGF-β and WNT pathways. Inhibition of the cross-activated pathways enhances the intensity of BMP4 signaling and its ability to induce the Totipotent state. Our study demonstrates that the cross-activation of multiple pathways by a single stimulus (e.g. BMP4) can lead to the induction of multiple cell fates, resulting in pronounced stem cell heterogeneity. We further demonstrate that such signaling complexity is amenable for reprogramming stem cell states.

In mouse preimplantation embryos, BMP4 signaling is observed to be active from the 4-Cell stage onward, is required for normal morula establishment, and regulates the establishment of extra-embryonic lineages and the preimplantation epiblast (*63, 64*). In agreement, the BMP4-driven induction of Totipotent cells in LBPXRS correlated better with the 4-Cell and 8-Cell stages (**Figure 6A-B**). Taken together with these observations, our study demonstrates that BMP4 signaling plays an essential role in establishing the Totipotent state. A recent study found that retinoic acid can also induce a 2C-like Totipotent state in ESCs (*27*). Although retinoic acid is not used to culture mouse ESCs as it induces their differentiation towards neuroectoderm lineage (*28*), it may act in conjunction with BMP4 signaling to induce the Totipotent state. Similar to previous reports, we also found that Totipotent cells existed in common culture conditions such as 2iL but were less apparent than in sL and LB (*17, 18*). How the WNT and LIF pathways activated in 2iL induce the Totipotent state in ESCs is not fully understood. It might involve WNT/LIF-mediated cross-activation of BMP4 or retinoic acid signaling, which remains to be characterized.

Our data reveal that BMP4 signaling is more complex than previously thought in ESCs. Surprisingly, it cross-activates multiple pathways required to promote alternative cell states. In addition, the intensity of BMP4 pathway activation itself is diminished by the cross-activations. Accordingly, BMP4 signaling could be enhanced by 2-3 fold when the cross-activations are inhibited (**Figure 4**). These data suggest that the BMP4 signaling information is prone to divergence for activating other pathways at the expense of its own activation. In addition to our findings for totipotency, BMP4 signaling has also been implicated in promoting pluripotency and extra-embryonic lineages of preimplantation embryos (*63, 64*). How BMP4 signaling regulates diverse cell types of early embryogenesis is not well understood. Our study suggests that tuning of BMP4 signaling activity combined with its ability to cross-activate multiple pathways may play an important role in orchestrating early embryogenesis.

How cell signaling influences stem cell heterogeneity is an active area of research. The FGF and TGF-β pathways can regulate cell state heterogeneity of mouse ESCs (*10, 11, 13*). The pluripotency genes Oct4 and Sox2 can induce FGF expression, resulting in the differentiation of ESCs (*29, 32*). The intensity of FGF pathway activation (i.e. ERK1/2) can regulate the transition of ESCs from the Pluripotent to the PrEn state (*65*). Similarly, TGF-β signaling regulates ESC heterogeneity (*66*). Our study reveals that BMP4 signaling plays an essential role in regulating ESC heterogeneity and demonstrates that a single stimulus’s cross-activation of multiple pathways is another crucial signaling parameter to explain ESC heterogeneity. While the parallel activation of multiple pathways by a stimulus signal is common, it is unclear whether this actually leads to meaningful response outcomes. Our data demonstrate that in the case of BMP4 signaling in ESCs, the cross-activation of the FGF, TGF-β and WNT pathways is not spurious but leads to the induction of their canonical target genes and the cell state genes (**Figures 2 & 4**). Signaling cross-activations could similarly regulate the heterogeneous cell fate specification commonly observed during developmental differentiation and cell reprogramming. Dissecting such mechanisms, as demonstrated, may provide new approaches to managing cellular heterogeneity in differentiation and reprogramming.

The totipotency of a cell is evaluated by its bidirectional developmental potential, and we show that Totipotent cells in LBPXRS possess this function (*17, 62*) (**Figures 6, S6 & S7**). At the transcriptome level, they resembled 4-Cell and 8-Cell stages more than the 2-Cell stage of preimplantation embryos (**Figure 6**). They also expressed multiple genes associated with the 2C-like state, suggesting that LBPXRS may induce a spectrum of Totipotent states instead of a single 2C-like state (**Figures 3 & 5**). Indeed, the Totipotent state is observed to be the property of cells from the Zygote to the 8-Cell stage of preimplantation embryos (*67–69*). Therefore, we referred to the reprogrammed cells in LBPXRS as “Totipotent” and not “2C-like” cells. Considering Totipotent cells from LBPXRS as *bona fide* 2C-like cells would require further characterization of zygotic genome activation signatures and the ability to reconstitute pre-and postimplantation embryos *in vitro* (*70*).

We observed about ~20% of ESCs reprogrammed to the Totipotent state in LBPXRS at the single-cell level (**Figures 3 & 5**). However, LBPXRS clearly induced the expression of Totipotent genes at the population level (**Figure S5C**). The refraction of other cells from conversion to the Totipotent state in LBPXRS potentially suggests alternative mechanisms as barriers. Potential barriers could be due to an incomplete suppression of histone modifiers (leading to a non-permissive chromatin state) and negative regulators of totipotency, as well as a partially reduced gene-splicing rate (*17, 71–73*). It may also be that additional signaling events such as retinoic acid are required to work synergistically with the activated BMP4 pathway to promote totipotency (*27*). Future studies oriented to address and overcome these limitations may result in a higher reprogramming efficiency of ESCs to the Totipotent state. A related future challenge is to design stable culture conditions for continuous propagation and expansion of exclusive Totipotent cells.

Even though our study reveals that BMP4 signaling cross-activates the FGF, TGF-β and WNT pathways, the precise molecular details of how these cross-activations are initiated or their exact paths of information processing remain to be established. These events could be triggered at the receptor level or through a specific network of intracellular signalling mediators. Our data is also limited in determining whether all the pathways are simultaneously activated at the single-cell level. Future systematic and quantitative studies of signal propagation using proteomics and reporter assays may reveal specific molecular details of cross-activation mechanisms.

## Acknowledgements

We are grateful to Alex Gregorieff, Nicole Francis and Gerardo Ferbeyre for helpful comments on the manuscript. We thank the Flow Cytometry, Molecular Biology and Functional Genomics, Microscopy and Imaging core facilities at the Montreal Clinical Research Institute (IRCM). We acknowledge the following funding support for this work: M.M. is supported by the Canadian Institutes of Health Research Project Grant 173273, Maud Menten New Principal Investigator Prize 173596, Fonds de Recherche Quebec - Santé Research Scholar Award 254497, and John R. Evans Leaders Fund from Canada Foundation for Innovation; T.M. is supported by the Emmanuel Triassi-IRCM Foundation Scholarship; V.H. is supported by the Scholarship Hommage Michel-Bélanger.

## Author Contributions

M.M. conceived and supervised the project. T.M., M.P., and V.H. designed and performed experiments and analyzed data. J.L. and E.L.d.R. performed single-cell RNA-seq data analysis. M.M. wrote the manuscript with inputs from all authors.

## Data availability

The high throughput single-cell RNA-seq data reported in this study are available on Gene Expression Omnibus under the accession number GSE198384.

**Supplementary Materials** include the following:

Figures S1–S7

Table S1

Data S1

## MATERIALS AND METHODS

### Cell culture

R1 (E14-Tg2A) mouse ESCs were routinely cultured by standard methods. Cells were cultured in knockout DMEM supplemented with non-essential amino acids, sodium pyruvate, L-Glutamine, β-mercaptoethanol, 15% fetal calf serum (Hyclone) and Leukemia inhibitory factor (LIF) (1,000 units/ml, Millipore ESG1106). Cell medium was replaced daily, and cells were passaged every two to three days. In LB and 2iL conditions, cells were cultured in N2B27 containing either LIF and BMP4 (10 ng/ml, Stemgent 03-0007) or CHIR99021 (3 μM, Stemgent 04-0004-02) and PD0325901 (1 μM, Millipore Sigma 444968) respectively.

### Immunofluorescence

Immunofluorescence was performed as previously described (*28*). ESCs were grown on tissue culture-treated plastic or coverslips in 24-well plates and fixed with 4% paraformaldehyde (PFA) in BBS (50 mM BES Sodium salt, 280 mM NaCl, 1.5 mM Na_2_KPO_4_ and pH 6.96) with 1 mM CaCl_2_ for 15 min. Cells were then washed thrice and blocked with BBT-BSA buffer (BBS with 0.5% BSA, 0.1% Triton and 1 mM CaCl_2_) for 45 min at room temperature. Cells were incubated in a humid chamber at 4°C overnight with primary antibodies, washed thrice with BBT-BSA then incubated with fluorophore-conjugated secondary antibodies for 1 to 2 h in the dark at room temperature. Cells were then washed with BBS-CaCl_2_, stained for DAPI, mounted using vectashield and imaged. Imaging was completed using a Nikon Ti motorized inverted microscope fitted with perfect focus and Yokagawa CSU-X1 spinning disk confocal systems, 20x Plan-Apochromatic objective (NA .75) and with a Hamamatsu ORCA-AG cooled CCD camera. A Lumencor SOLA fluorescence light source and a QUAD 405/491/561/642 dichroic mirror filters were used for excitation and emission, respectively. Images were acquired using MetaMorph software and analyzed using Fiji (ImageJ). Primary antibodies and their dilutions were used as follows: anti-OCT4 (Santa Cruz sc-5279, 1:300); anti-NANOG (eBioscience 14-5761, 1:300); anti-MERVL-gag (Epigentek A-2801, 1:100); anti-Brachyury (T) (Santa Cruz sc-17743, 1:300); anti-β-Catenin (eBioscience 14-2567-82, 1:150) and anti-Gata6 (Santa Cruz sc-7244, 1:500). Secondary antibodies conjugated to either Alexa Fluor 488, 568 or 647 fluorophores (ThermoFisher Scientific) were used at a 1:500 dilution.

### Flow cytometry

Cells were harvested, fixed with 4% PFA for 15 min, then washed with 1% BSA in phosphate buffered saline (PBS). Fixed cells were permeabilized with cold 80% EtOH for 15 min, then washed twice with 1% BSA in PBS. Cells were stained for 60 min and then washed twice with 1% BSA in PBS. Cells were stained against OCT4, MERVL-gag and NANOG with primary antibodies and Alexa Fluor conjugated secondary antibodies as previously described above for immunofluorescence. Following antibody staining, cells were filtered to remove aggregates, and fluorescence data were collected on the LSRII (BD Biosciences) flow cytometer equipped with 405 nm, 488 nm, 561 nm, and 633 nm lasers. Flow cytometry data were analyzed using FlowJo and Cytobank.

### Single-cell RNA sequencing (scRNA-seq)

Cells grown in either LB or LBPXRS for 72 h were collected, kept on ice throughout preparations, and then passed through a 40 μM strainer. Single-cell suspensions were prepared with cell viability >90% in PBS with 0.04% non-acetylated BSA, then loaded onto a Next GEM Chip A (PN-1000009) with the Chromium Single Cell 3’ Gel Bead Kit v2 (PN-120235) as described by the manufacturer. Single cells were captured onto a 10X Genomics Chromium controller with a recovery target of 5000 cells per sample.

#### Library construction

cDNA was generated following instructions provided by 10X Genomics. Size selection and quality assessment were completed using Ampure XP beads (Beckman Coulter A63881) and the Bioanalyzer High Sensitivity DNA Kit (Agilent 5067-4626). Next, ¼ of total cDNA was used to generate libraries using the Chromium Single Cell 3’ Library Kit v2 (PN-120234) and barcoded using the Chromium Multiplex Kit (PN-120262) following instructions provided by the manufacturer. Libraries were then size selected, and size distribution was assessed using the Bioanalyzer High Sensitivity DNA Kit with an average size of 470 bp. The library was sequenced on the Illumina Novaseq S Prime flowcell with a depth of 57,000-94,000 PE28-91 reads per cell.

### Conditions to manipulate signaling cross-activation

For all the ligands and small molecules inhibitors tested, cells were cultured in an N2B27 medium supplemented with LIF (1,000 units/ml), non-essential amino acids, L-Glutamine and β-mercaptoethanol. ES cells cultured in sL were dissociated, seeded in the indicated conditions, cultured for 72 h with a daily change of medium, then collected for analysis (western blot or flow cytometry). For kinetics with cycloheximide, cells were seeded and allowed to attach for 24 h in sL. Then the sL medium was removed, cells were washed twice with PBS, LB containing cycloheximide was added at t=0 and cells were harvested at indicated time points for western blot analysis. Except where indicated otherwise, the ligands and small molecule inhibitors used in this study and their concentration were: LIF (1,000 units/ml, Millipore ESG1106), BMP4 (10 ng/ml, Stemgent 03-0007), CHIR99021 (3 μM, Stemgent 04-0004-02), PD0325901 (1 μM, Millipore Sigma 444968), XAV939 (10 μM, Millipore Sigma 575545), TAK1 inhibitor (1 μM, Tocris 3604), SB431542 (2 μM, Millipore Sigma 616464), LDN193189 (500 nM, Sigma Aldrich SML0559), STAT3 inhibitor VI (100μM, Calbiochem 573102) and cycloheximide (100 mg/ml, Sigma 01810). All compounds were reconstituted and used as recommended by the manufacturer.

### Western blots

Cells were rinsed with PBS and lysed in Laemmli 2x buffer (4% SDS, 20% glycerol, 120 mM Tris-HCl pH 6.8). Extracts were boiled for 10 min, and the protein concentration was assessed with NanoDrop (ThermoFisher). Samples were supplemented with β-mercaptoethanol and bromophenol blue. Extracts were loaded onto SDS-PAGE gels and then transferred to polyvinylidene difluoride (PVDF) membranes. Membranes were incubated with the following primary antibodies: anti-GAPDH (6C5) (Invitrogen AM4300) lot 00755156, anti-phosphoSMAD1/5 (S463/465) (41D10) (Cell Signaling 9516) lot 9, anti-SMAD1 (D59D7) (Cell Signaling 6944T) lot 2, anti-phosphoERK1/2 (T202) (D13.14.4E) (Cell Signaling 4370S) lot 24, anti-ERK1/2 (137F5) (Cell Signaling 4695S) lot 28, anti-β-Catenin (D10A8) (Cell Signaling 8480S) lot 5, anti-phospho-SMAD2 (S465/467)/SMAD3 (S423/425) (D27F4) (Cell Signaling 8828S) lot8 and anti-SMAD2/3 (Santa Cruz sc-133098) lot L0820. Membranes were imaged using ImageQuant LAS 4000 (GE Healthcare). Bands were quantified from immunoblot pictures using Fiji (ImageJ). Signal intensities were normalized over GAPDH intensity.

### RT-qPCR

Total RNA was purified from samples using the standard TRIzol method. Complementary DNA (cDNA) was prepared using the 5X Prime Script RT Master Mix (Takara RR-036A-1) with 1 μg of total RNA. Reverse transcription was carried out with the standard cycling condition as per the manufacturer’s instructions. cDNA was diluted 30-fold, and quantitative PCR (qPCR) was performed using the PowerUp SYBR Green Master Mix (Thermo Fisher Scientific 100029283). The relative gene expression fold change was calculated using the ΔΔCt method and plotted based on the log-fold expression. The primers used for qPCR are listed in Table S1.

### Chimera formation

ESCs containing MERVL::tdTomato were infected with lentivirus encoding EF1α::EGFP to sort Totipotent cells and track the blastocyst embryo complementation of microinjected cells. Cells constitutively and stably expressing EGFP under the EF1α promoter were sorted, cultured in LBPXRS then sorted by flow cytometry to obtain Pluripotent (tdTomato-EGFP+) and Totipotent (tdTomato+ EGFP+) cells.

#### Blastocyst chimera formation

The corresponding sorted, single Pluripotent or Totipotent cells were microinjected into the E2.5 morulas obtained from C57BL/6N mice. Microinjected morulas were incubated for 24 h at 37ΰC, followed by their imaging using a fluorescence microscope. 87 and 98 morula embryos were microinjected with individual Pluripotent cells and Totipotent cells, respectively.

#### Embryo chimera formation

Sorted Pluripotent or Totipotent cells were microinjected into E2.5 C57BL/6N morulas, implanted into pseudopregnant females the following day and allowed to develop until E12.5 day embryos. The mature embryos were collected from chimeric mice at E12.5.

### Immunofluorescence staining of E12.5 chimeras

Mature embryos were collected from chimeric mice at E12.5, fixed for 2 h with 4% PFA, washed overnight with PBS, incubated for 4 h in 30% sucrose, and frozen in OCT. Multiple 10 μm longitudinal cryosections were taken from the mid-embryo and frozen at −80°C. On the day of staining, sections were let to reach room temperature and were first processed for antigen retrieval using 10 mM citrate buffer (pH 6.0) at 100°C for 10 min, then cooled to room temperature. Sections were then permeabilized with 0.3% Triton X-100 in PBS and blocked with PBS with 1% BSA, 10% FBS and 0.3% Triton X-100 for 1 h at room temperature. Overnight staining was performed at 4°C with the following primary antibodies: Cdx2 (1:150), Sox1 (1:300), Brachyury (1:300) and Tpbpa (1:100) diluted in PBS with 1% BSA, 1% FBS and 0.2% Triton X-100. The following day, sections were stained with the appropriate secondary antibodies for 1 h at room temperature, washed with PBS, counterstained with 4,6-diamidino-2-phenylindole and imaged with a LEICA SP8 confocal microscope. Acquired images were analyzed using Fiji (ImageJ).

### scRNA-seq analysis

Single-cell matrices were generated using Cellranger (v.4.0.0) provided by 10X genomics. Sequences were aligned with the mm10 mouse transcriptome. Matrices were imported and analyzed with the R package “Seurat” (v.4.0.3) (*56*). We filtered out genes expressed in less than 3 cells and low-quality cells expressing less than 200 genes. Cells with more than 50 000 transcripts were removed as they likely represent cell doublets. Cells with more than 10% of genes from the mitochondrial genome were also removed as dying cells often exhibit extensive mitochondrial contamination. Cells that did not express any cell state markers were excluded from the analysis. Each matrix was normalized with Seurat’s “NormalizeData” function using “LogNormalize” as the normalization method and a scale factor equal to 10 000 (default value). Matrices were integrated using the “IntegrateData” function. Linear dimensionality reduction was performed using the “RunPCA” function. Non-linear dimensionality reduction was then performed using the “RunUMAP” function, considering the Euclidean distances of the first 30 PCA components. Plots showing gene expression value on the UMAP space **(Figure S4)** were generated using the “FeaturePlot” function. kNN graph construction was performed using the “FindNeighbors” function, which also considered the first 30 PCA components. Clustering (Louvain method) was performed using the “FindClusters” function, with a resolution equal to 0.8. Differential expression analysis was performed using the “FindAllMarkers” function to detect markers considering positive log-transformed fold change values above 0.25. Finally, trajectory analysis **(Figure 5D-E)** was performed with the R package “Monocle3” (v.0.2.3.0) (*60*) using totipotent cells as the source of the pseudotime variable (pseudotime = 0). To display trajectory analysis results, UMAP was performed on the LB/LBPXRS integrated scRNAseq matrix using the “reduce_dimension” function found in the Monocle3 package.

#### Integration of scRNA-seq data

The integration of Totipotent cells with preimplantation cell stages and *in vitro* generated 2C-like cells was performed through Scanpy’s BBKNN integration pipeline. To perform this analysis, we integrated Totipotent state cells in LBPXRS (greater than 95^th^ percentile Gata2 expression) with multiple scRNA-seq datasets: Zygote, 2-Cell (early, mid and late), 4-Cell, 8-Cell and Morula from Deng et.al. (*74*); 1-Cell and 2-Cell from Huang et.al. (*57*); E3.0-3.5 and E4.5 from Mohammed et.al. (*75*); Primed cells from Chen et.al. (*76*); 2i cells from Chen et.al. (*76*) and Posfai et.al. (*70*) (GSE145609); and the in-vitro generated expanded pluripotent stem cells (EPSCs) from Yang et.al. (*25*) (EPSC-1) and Yang et.al. (*26*) (EPSC-2). The Morula cells consisted of both 16-Cell and E2.5-2.75 stage cells. A correlogram plot was generated with Scanpy’s “correlation_matrix” function considering the highly variable genes deducted from Scanpy’s “highly_variable_genes” function.

## SUPPLEMENTARY FIGURE LEGENDS

**Figure S1:**
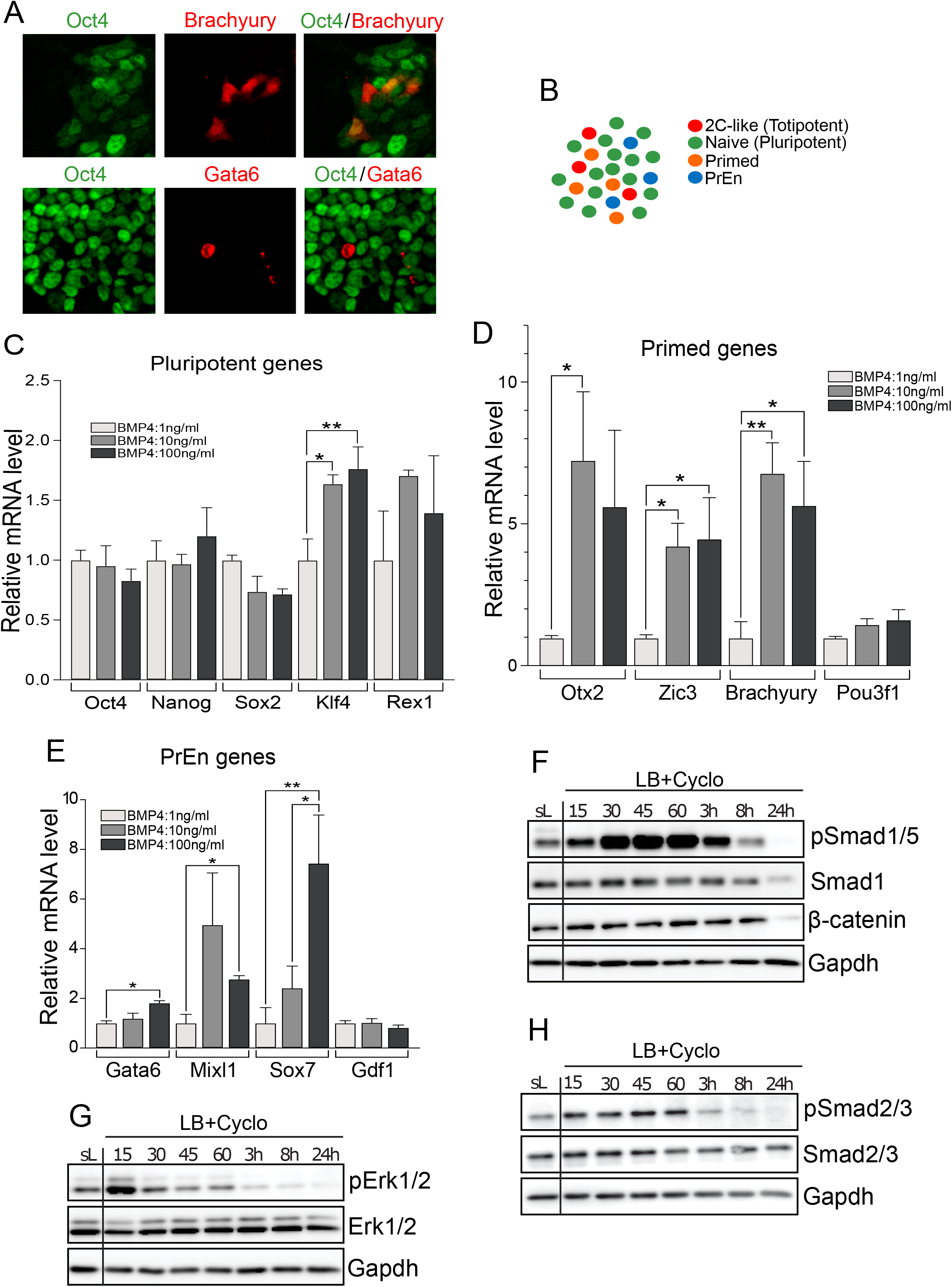
BMP4 induces multiple cell states in ESCs. **(A)** Immunofluorescence images of Oct4 (green) and Brachyury (red) or Gata6 (red) stained ESCs cultured in sL. **(B)** Schematic showing multiple cell states observed in cultured mouse ESCs: Totipotent (2C-like), Pluripotent (Naive), Primed and PrEn. Relative expression of the indicated Pluripotent **(C)**, Primed **(D)**, and PrEn **(E)** genes in mouse ESCs cultured in LB for 72 h. Error bars indicate the mean ± SEM of four biological replicates. ANOVA: *p≤ 0.05; **p≤ 0.01. Western blot analysis of BMP4 and WNT **(F)**, FGF **(G)**, TGF-β **(H)** signaling pathway activation in ESCs cultured in LB in the presence of cycloheximide (Cyclo) at 100 μg/ml for different time points (15 min, 30 min, 45 min, 60 min, 3 h, 6 h and 24 h). Cells cultured for 15 min in sL with Cyclo at 100 μg/ml were used as a control. Gapdh was used as a loading control, and the displayed images are representative of four independent experiments.

**Figure S2:**
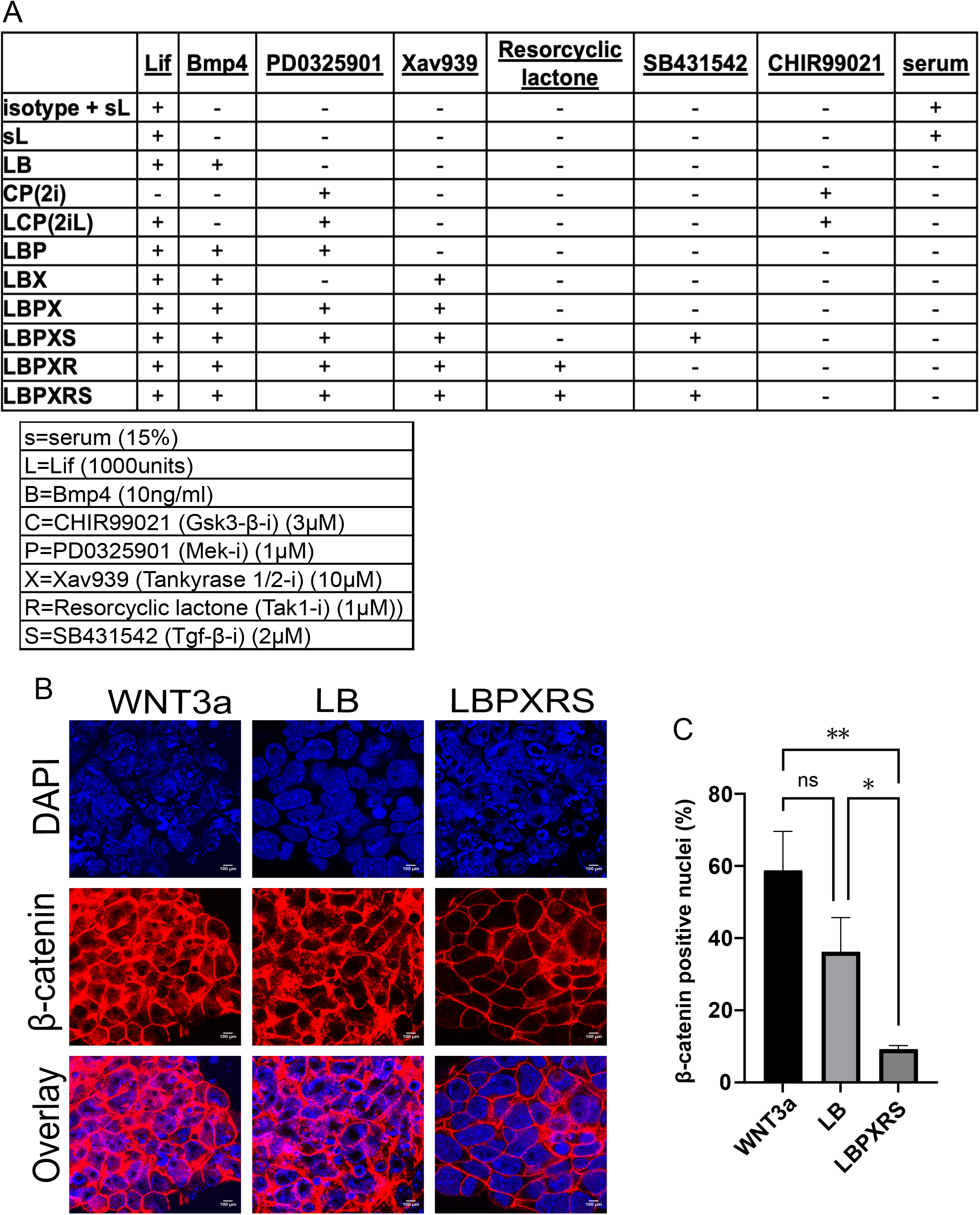
Rational inhibition of the BMP4 cross-activated FGF, TGF-β and WNT pathways. **(A)** Combinations of ligands and/or small molecule inhibitors used to either activate or inhibit a signaling pathway. The concentration and the expanded name of the acronym used for each ligand or inhibitor are listed at the bottom. “+” and “-” indicates the addition or exclusion of the corresponding ligand or inhibitor in the basal N2B27 medium. **(B)** Immunostaining of β-catenin (red) and DAPI staining (blue) of ESCs cultured in LB, LBPXRS or treated with WNT3a. Scale bar, 100 μm. **(C)** The proportion of cells with nuclear β-catenin in the indicated conditions. ANOVA: ns, not-significant; *p≤ 0.05, **p≤ 0.015.

**Figure S3:**
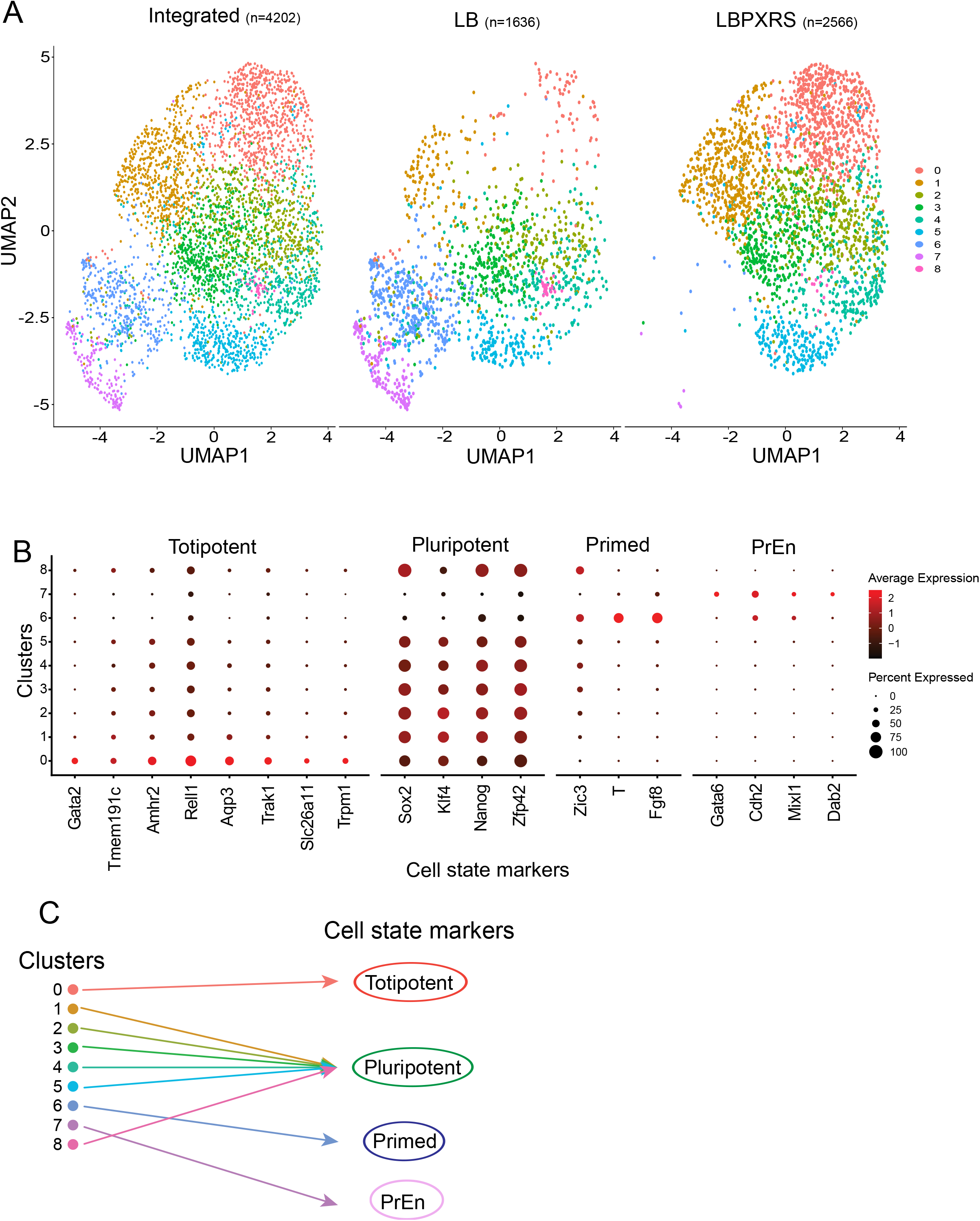
Integration of scRNA-seq datasets of ESCs cultured in LB and LBPXRS. **(A)** UMAP space showing Louvain clusters in LB or LBPXRS and their integrated dataset. Cells were partitioned into clusters using the Louvain community-detection algorithm. **(B)** Cell proportions expressing Totipotent, Pluripotent, Primed and PrEn genes and their average expression in each cluster. **(C)** Schematic showing the annotation of Louvain clusters to either Totipotent, Pluripotent, Primed and PrEn cell type based on the cell state genes expression observed in B and Figure S4.

**Figure S4:**
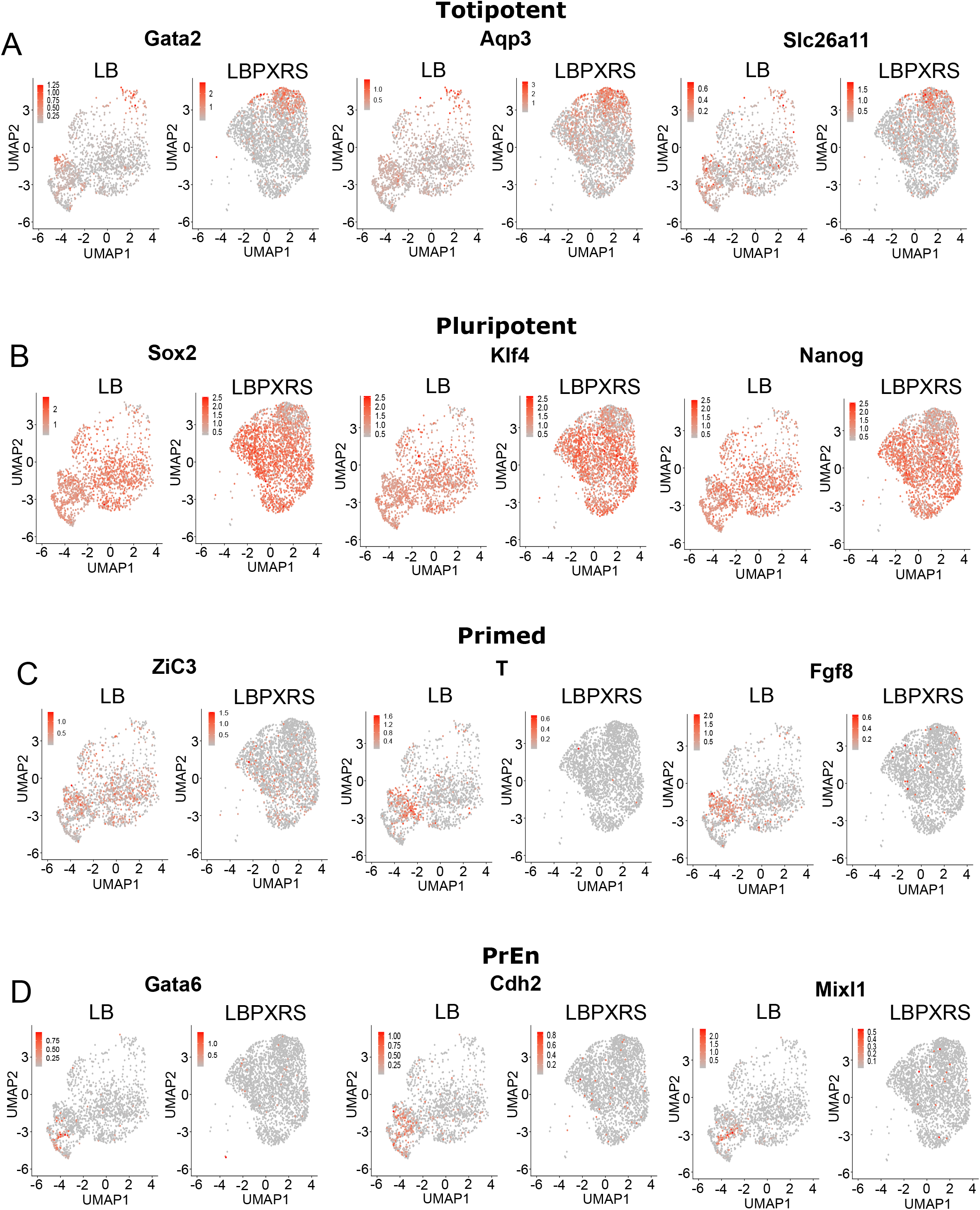
Cell state gene expression levels in ESCs cultured in LB and LBPXRS. The relative expression level of Totipotent **(A)**, Pluripotent **(B)**, Primed **(C)** and PrEn **(D)** cell state genes in ESCs cultured in LB and LBPXRS.

**Figure S5:**
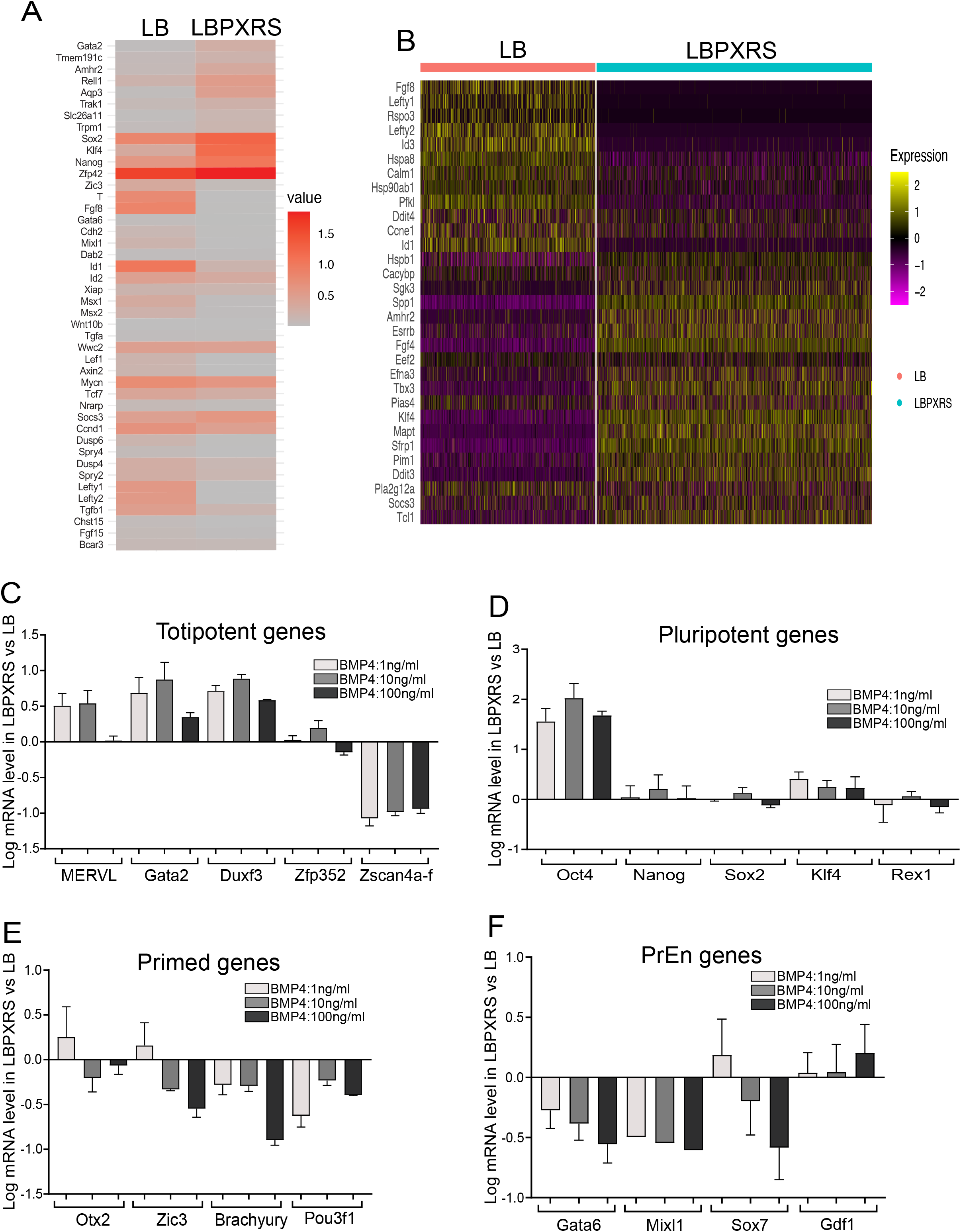
Differential expression of cell state and signaling genes in LB and LBPXRS. **(A)** Average expression heatmap of Totipotent, Pluripotent, Primed and PrEn cell state genes and common signaling genes. **(B)** Heatmap of top differentially expressed genes between LB and LBPXRS. Relative expression (log scale, relative to LB) of Totipotent **(C)**, Pluripotent **(D)**, Primed **(E)** and PrEn **(F)** state genes in ESCs with increasing concentrations of BMP4. Error bars indicate the mean ± SEM of four to eight biological replicates.

**Figure S6:**
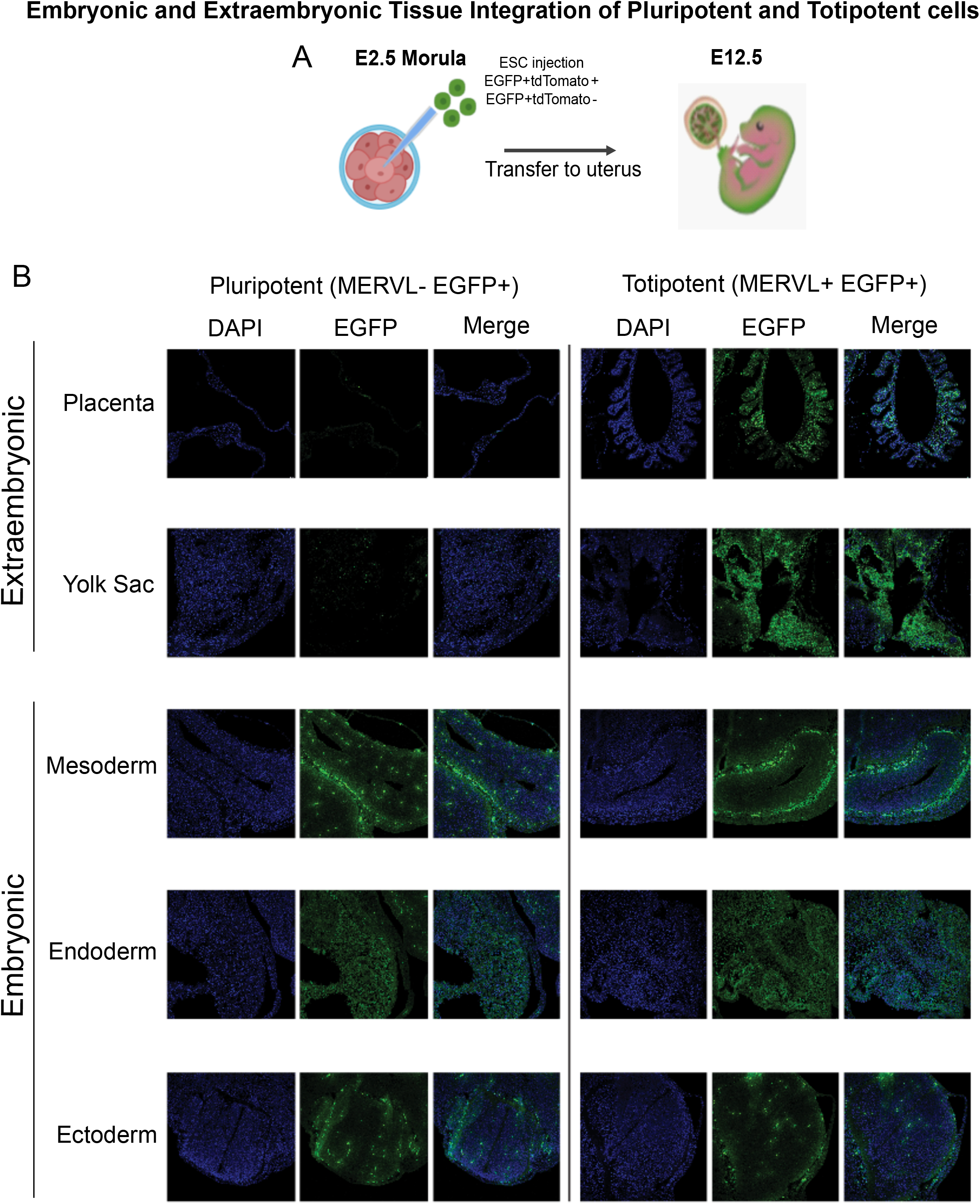
Embryonic and extra-embryonic tissue integration of Pluripotent and Totipotent cells from LBPXRS. **(A)** Schematic showing the postimplantation embryo chimera generation procedure using microinjection of EGFP-labeled cells into the morula of the preimplantation embryo. **(B)** Cryosections of embryonic and extra-embryonic tissues from E12.5 embryos showing the integration of sorted and microinjected Pluripotent (MERVL-EGFP+) and Totipotent (MERVL+ EGFP+) cells from LBPXRS. Sections were imaged for EGFP (green) and nuclear DNA using DAPI staining (blue). Overlay images of DAPI and EGFP are also shown.

**Figure S7:**
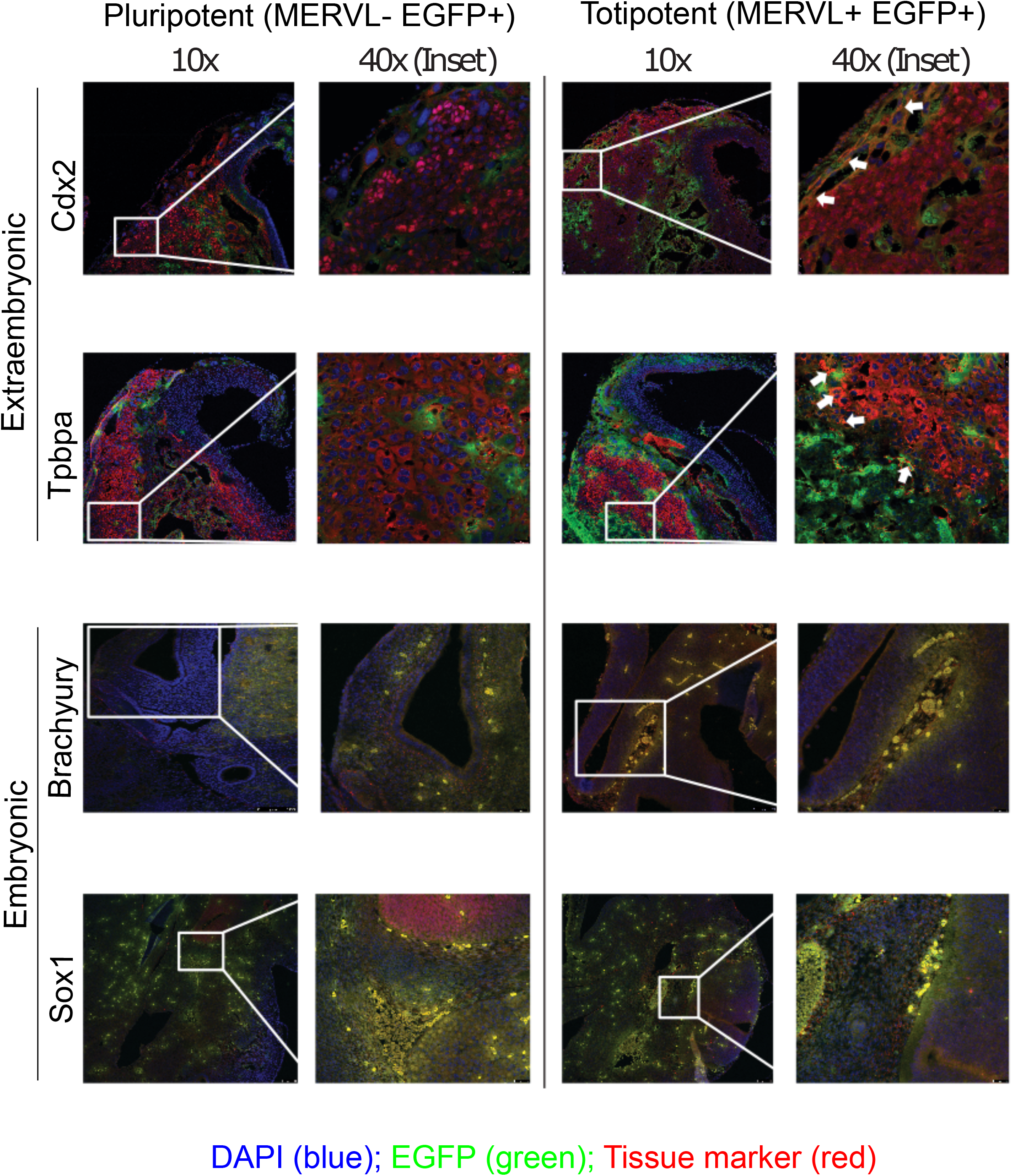
Evaluation of *in vivo* tissue integration of Pluripotent and Totipotent cells from LBPXRS. Immunohistochemistry of the cryosections of E12.5 chimeric embryos generated using Pluripotent (MERVL-EGFP+) and Totipotent (MERVL+ EGFP+) cells from LBPXRS condition. The cryosections were stained for embryonic tissue-specific markers (Brachyury for Mesendoderm; Sox1 for Neuroectoderm) and extra-embryonic tissue-specific markers (Cdx2 and Tpbpa), and nuclear DNA using DAPI. The stained cryosections were simultaneously imaged for the tissue marker (red) along with EGFP (green) and DAPI (blue). The left panel shows a wider view of the tissue section, scale bar: 100 μm. The right panel shows a magnified plane of a selected section of the image, scale bar: 25 μm. The white arrows are used to highlight the cells with co-localized signals of the tissue marker and EGFP.

